# Experimentally-validated correlation analysis reveals new anaerobic methane oxidation partnerships with consortium-level heterogeneity in diazotrophy

**DOI:** 10.1101/2020.04.12.038331

**Authors:** Kyle S. Metcalfe, Ranjani Murali, Sean W. Mullin, Stephanie A. Connon, Victoria J. Orphan

**Affiliations:** Division of Geological and Planetary Sciences, California Institute of Technology, Pasadena, CA USA

## Abstract

Archaeal anaerobic methanotrophs (‘ANME’) and sulfate-reducing Deltaproteobacteria (‘SRB’) form symbiotic multicellular consortia capable of anaerobic methane oxidation (AOM), and in so doing modulate methane flux from marine sediments. The specificity with which ANME associate with particular SRB partners *in situ*, however, is poorly understood. To characterize partnership specificity in ANME-SRB consortia, we applied the correlation inference technique SparCC to 310 16S rRNA Illumina iTag amplicon libraries prepared from Costa Rica sediment samples, uncovering a strong positive correlation between ANME-2b and members of a clade of Deltaproteobacteria we termed SEEP-SRB1g. We confirmed this association by examining 16S rRNA diversity in individual ANME-SRB consortia sorted using flow cytometry and by imaging ANME-SRB consortia with fluorescence *in situ* hybridization (FISH) microscopy using newly-designed probes targeting the SEEP-SRB1g clade. Analysis of genome bins belonging to SEEP-SRB1g revealed the presence of a complete *nifHDK* operon required for diazotrophy, unusual in published genomes of ANME-associated SRB. Active expression of *nifH* in SEEP-SRB1g and diazotrophic activity within ANME-2b/SEEP-SRB1g consortia was then demonstrated by microscopy using hybridization chain-reaction (HCR-) FISH targeting *nifH* transcripts and by FISH-nanoSIMS experiments. NanoSIMS analysis of ANME-2b/SEEP-SRB1g consortia incubated with a headspace containing CH_4_ and ^15^N_2_ revealed differences in cellular ^15^N-enrichment between the two partners that varied between individual consortia, with SEEP-SRB1g cells enriched in ^15^N relative to ANME-2b in one consortium and the opposite pattern observed in others, indicating both ANME-2b and SEEP-SRB1g are capable of nitrogen fixation, but with consortium-specific variation in whether the archaea or bacterial partner is the dominant diazotroph.

## Introduction

The partnership between anaerobic, methanotrophic Archaea (ANME) and their associated sulfate-reducing bacteria (SRB) is one of the most biogeochemically-important symbioses in the deep-sea methane cycle [1, 2]. As a critical component of methane seep ecosystems, multicellular consortia of ANME and associated SRB consume a significant fraction of the methane produced in marine sediments, using sulfate as a terminal electron acceptor to perform the anaerobic oxidation of methane (AOM) [1–4]. ANME-SRB consortia are thought to perform AOM through the direct extracellular transfer of electrons between ANME and SRB [5–7]. Along with symbiotic extracellular electron transfer, ANME-SRB consortia also exhibit other traits of mutualism such as the sharing of nutrients. For example, members of the ANME-2 clade have been reported to fix and share N with partner bacteria [8–11], but the extent to which diazotrophic capability might vary across the diverse clades of ANME and associated SRB is the focus of ongoing research.

Comparative studies of ANME [12] and associated SRB [13, 14] genomes from multiple ANME-SRB consortia have revealed significant diversity across clades, particularly for SRB genomes falling within subclades of the SEEP-SRB1 [14], common SRB partners to ANME [15]. However, the implications of symbiont diversity for metabolic adaptation in ANME-SRB consortia are obscured by the absence of clearly-established ANME-SRB pairings in the environment. A framework defining these pairings would address this gap in knowledge. Establishing this framework for partnership specificity in ANME-SRB consortia—being the preference that certain ANME exhibit for specific SRB partners—would shed light on the extent to which ANME or SRB physiology may differ in consortia constituted of different ANME-SRB pairs.

As an aspect of ANME or SRB physiology that may differ in different ANME-SRB pairings, nitrogen anabolism has been observed to be involved in the symbiotic relationship between partners [8, 9] and has been shown to influence niche differentiation of different ANME-SRB consortia via nitrate assimilation ability [16]. Previous evidence documenting active diazotrophy in ANME-SRB consortia by nitrogenase expression [8] and ^15^N_2_ fixation by nanoSIMS indicated that ANME-2 are the primary diazotrophs in ANME-SRB consortia and supply fixed nitrogen to SRB partners [8–10]. The diazotrophic potential of syntrophic SRB, however, and their role in nitrogen fixation within consortia is poorly understood. Evidence from SRB genomes [14] and the expression of unidentified nitrogenase sequences in methane seep sediments [8] suggested that seep associated SRB may fix nitrogen, opening up the possibility of variation in diazotrophic activity among taxonomically-distinct ANME-SRB consortia.

Previous research characterizing the diversity of partnerships in ANME-SRB consortia have employed fluorescence microscopy, magnetic separation by magneto-FISH, and single-cell sorting techniques (e.g. BONCAT-FACS) that are robust against false positives, but are often limited in statistical power. Fluorescence *in situ* hybridization (FISH) has helped to establish the diversity of ANME-bacterial associations, with ANME constituting four diverse polyphyletic clades within the Methanomicrobia: ANME-1a/b [4, 17–20], ANME-2a,b,c [3, 20–22], ANME-2d [23, 24], and ANME-3 [20, 25, 26]. ANME-associated SRB have also observed by FISH to be diverse, representing several clades of Deltaproteobacteria including the *Desulfococcus/Desulfosarcina* (DSS) clade [3–6, 15, 19–22, 27–33], two separate subclades within the Desulfobulbaceae [16, 25, 26], a deeply-branching group termed the SEEP-SRB2 [34], and a thermophilic clade of Desulfobacteraceae known as HotSeep-1 [34, 35]. These FISH studies documented associations for different ANME-SRB consortia, including partnerships between members of ANME-1 and SEEP-SRB2 [13] or HotSeep-1 [7, 13, 35], ANME-2a and SEEP-SRB1a [15], ANME-2c and SEEP-SRB1a [5], SEEP-SRB2 [13, 34], or Desulfobulbaceae [29], and ANME-3 and SEEP-SRB1a [15] or Desulfobulbaceae [25, 26]. Conspicuously, SRB found in consortia with ANME-2b have only been identified broadly as members of the Deltaproteobacteria targeted by the probe S-C-dProt-0495-a-A-18 (often referred to as Δ495) [5, 31, 36], leaving little known about the specific identity of this SRB partner. Visualizing ANME-SRB partnerships by FISH has been a valuable aspect of AOM research, but FISH requires the design of probes with sufficient specificity to identify partner organisms and thus will only detect partnerships consisting of taxa for which phylogenetic information is known [22]. Magneto-FISH [29, 37, 38] or BONCAT-enabled fluorescence-activated cell sorting (BONCAT-FACS) of single ANME-SRB consortia [39] complement FISH experiments by physical capture (via magnetic beads or flow cytometry, respectively) and sequencing of ANME and associated SRB from sediment samples. These studies corroborated some of the patterns observed from FISH experiments, showing associations between ANME-2 and diverse members of the DSS [39]. Magneto-FISH and BONCAT-FACS observations of ANME-SRB pairings are also highly robust against false positives but can lack the statistical power conferred by more high-throughput approaches that is necessary to establish a general framework for partnership specificity.

Recently, a number of correlation analysis techniques have been introduced in molecular microbial ecology studies, providing information about patterns of co-occurrence between 16S rRNA OTUs or ASVs recovered from environmental iTag [40] diversity surveys [41–43]. Correlation analysis performed on 16S rRNA amplicon surveys provides a complementary method to Magneto-FISH and/or BONCAT-FACS that can be used to develop hypotheses about potential microbial interactions. While predictions of co-occurrence between phylotypes from these correlation analysis techniques have been reported in a number of diverse environments, they are rarely validated through independent approaches, with a few notable exceptions [44].

Here, we present a framework for ANME-SRB partnership specificity, using correlation analysis of 16S iTag amplicon sequences from a large-scale survey of seafloor methane seep sediments near Costa Rica to predict potential ANME-SRB partnerships. A partnership between ANME-2b and members of an SRB group previously not known to associate with ANME (SEEP-SRB1g) was hypothesized by correlation analysis and independently assessed using FISH and amplicon data from BONCAT-FACS-sorted ANME-SRB consortia. With this new framework, we were able to identify a novel partnership between ANME-2b and SEEP-SRB1g and map predicted physiological traits of SEEP-SRB1g genomes onto partnership specificity with ANME-2b. Our approach led us to formulate new hypotheses regarding how SEEP-SRB1g physiology may complement ANME-2b physiology, focusing on nitrogen fixation in SEEP-SRB1g. We demonstrate in this study that the symbiotic relationship between ANME and associated SRB can vary depending on the nature of the partner taxa and affirm the importance of characterizing individual symbiont pairings in understanding AOM symbiosis.

## Materials and Methods

Here, we present an abridged description of the methods used in this study. A full description can be found in the Supplemental Materials and Methods.

### Sample origin and processing

Pushcore samples of seafloor sediment were collected by DSV *Alvin* during the May 20-June 11 2017 ROC HITS Expedition (AT37-13) aboard R/V *Atlantis* to methane seep sites southwest of Costa Rica [45–47]. After retrieval from the seafloor, sediment pushcores were extruded aboard R/V *Atlantis* and sectioned at 1-3 cm intervals for geochemistry and microbiological sampling using published protocols [21, 48]. Samples for DNA extraction were immediately frozen in liquid N_2_ and stored at −80°C. Samples for microscopy were fixed in 2% paraformaldehyde for 24 h at 4°C. A full list of samples used in this study can be found in Supplemental Table 1 and additional location and geochemical data can be found at https://www.bco-dmo.org/dataset/715706.

### DNA extraction and iTag sequencing

DNA was extracted from 310 samples of Costa Rican methane seep sediments and seep carbonates (Supp. Table 1) using the Qiagen PowerSoil DNA Isolation Kit 12888 following manufacturer directions modified for sediment and carbonate samples [21, 49]. The V4-V5 region of the 16S rRNA gene was amplified using archaeal/bacterial primers, 515F (5’- GTGYCAGCMGCCGCGGTAA-3’) and 926R (5’-CCGYCAATTYMTTTRAGTTT-3’) with Illumina adapters [50]. PCR reaction mix was set up in duplicate for each sample with New England Biolabs Q5 Hot Start High-Fidelity 2x Master Mix in a 15 μL reaction volume with annealing conditions of 54°C for 30 cycles. Duplicate PCR samples were then pooled and 2.5 μL of each product was barcoded with Illumina NexteraXT index 2 Primers that include unique 8-bp barcodes. Amplification with barcoded primers used annealing conditions of 66°C and 10 cycles. Barcoded samples were combined into a single tube and purified with Qiagen PCR Purification Kit 28104 before submission to Laragen (Culver City, CA, USA) for 2 x 250 bp paired end analysis on Illumina’s MiSeq platform. Sequence data was submitted to the NCBI Sequence Read Archive as Bioproject PRJNA623020. Sequence data was processed in QIIME version 1.8.0 [51] following Mason, et al. 2015 [52]. Sequences were clustered into *de novo* operational taxonomic units (OTUs) with 99% similarity [53], and taxonomy was assigned using the SILVA 119 database [54]. The produced table of OTUs detected in the 310 methane seep sediment and seep carbonate amplicon libraries was analyzed using the correlation algorithm SparCC [41]. To examine phylogenetic placement of SRB 16S rRNA gene amplicon sequences predicted by network analysis to associate with particular ANME subgroup amplicon sequences, a phylogeny was constructed using RAxML-HPC [55] on XSEDE [56] using the CIPRES Science Gateway [57] from full-length 16S rRNA sequences of Deltaproteobacteria aligned by MUSCLE [58]. Genomes downloaded from the IMG/M database were searched using tblastn. Chlorophyllide reductase BchX (WP011566468) was used as a reference sequence for a tblastn *nifH* search using BLAST+. Genome trees were constructed using the Anvi’o platform [59] using HMM profiles from a subset [60] of ribosomal protein sequences and visualized in iTOL [61].

### FISH probe design and microscopy

A new FISH probe was designed in ARB [62]. This new probe, hereafter referred to as Seep1g-1443 (Supp. Table 2), was designed to complement and target 16S rRNA sequences in a monophyletic “*Desulfococcus* sp.” clade. Based on phylogenetic analysis (see below), this clade was renamed SEEP-SRB1g. Seep1g-1443 was ordered from Integrated DNA Technologies (Coralville, IA, USA). FISH reaction conditions were optimized for Seep1g-1443, with optimal formamide stringency found to be 35% (Supp. Fig. 1). FISH and hybridization chain reaction (HCR-) FISH was performed on fixed ANME-SRB consortia using previously published density separation and FISH protocols [22]. FISH was performed overnight (18 hr) using modifications (G. Chadwick, pers. comm.) to previously-published protocols [29, 39, 63, 64]. Structured-illumination microscopy (SIM) was performed on FISH and HCR-FISH (see below) experiments to image ANME-SRB consortia using the Elyra PS.1 SIM platform (Zeiss, Germany) and an alpha Plan-APOCHROMAT 100X/1.46 Oil DIC M27 objective. Zen Black software (Zeiss) was used to construct final images from the structured-illumination data.

**Figure 1.**
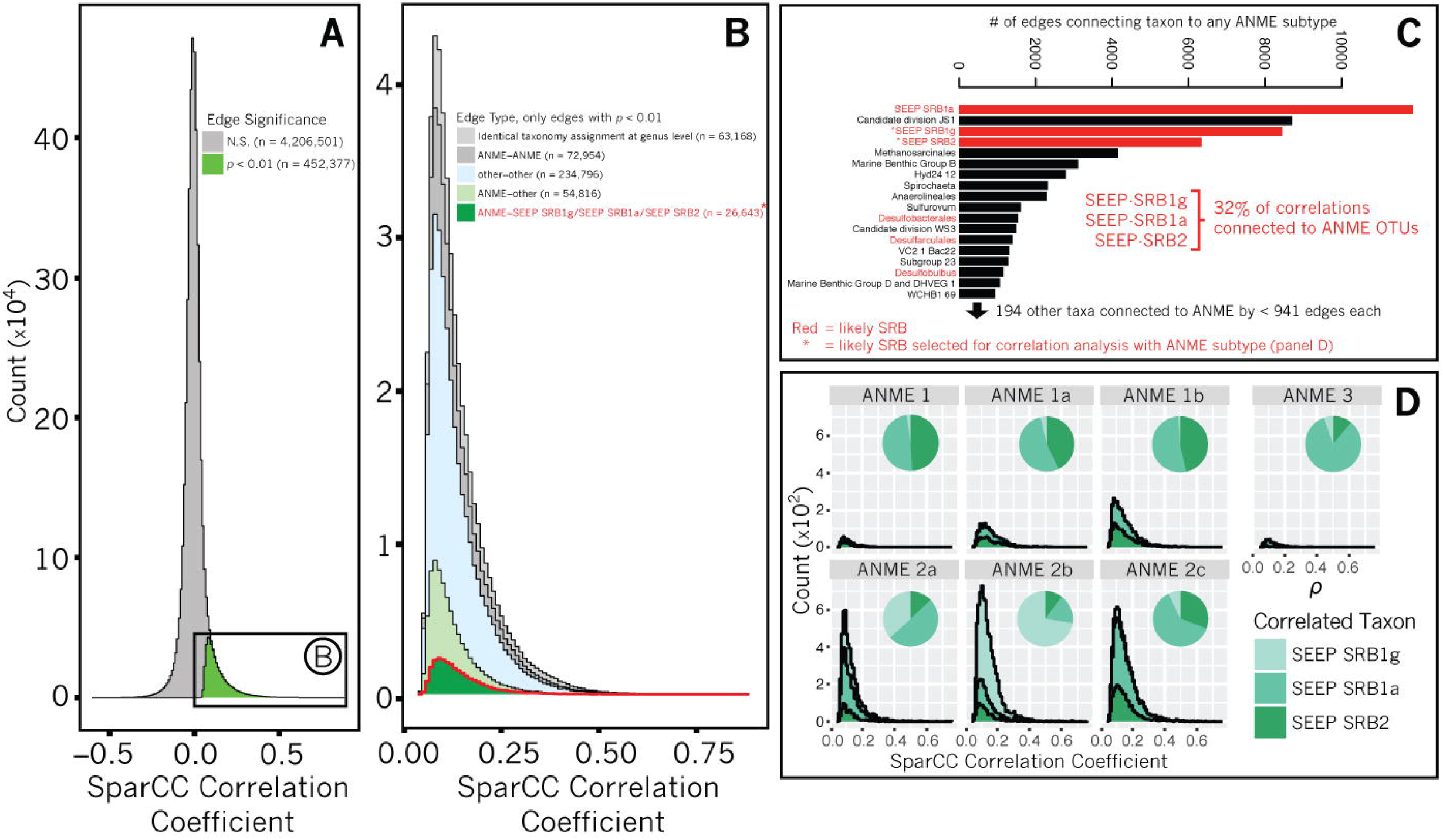
Analysis of SparCC-calculated correlations between 16S iTag amplicon sequences (OTUs clustered at 99% similarity) from an ecological survey of 310 methane seep sediment samples from seafloor sites off of Costa Rica. A stacked histogram (A) illustrates the proportion of correlations deemed significant on the basis of pseudo-*p*-values < 0.01 calculated by comparison with 100 bootstrapped correlation tables (see Materials and Methods). Of the correlations with pseudo-*p*-values < 0.01, 18% include ANME with a non-ANME taxon (B). Significant correlations between OTUs with taxonomy assignments that are identical at the genus level (e.g. two Anaerolinea OTUs) are indicated by identical taxonomy assignment. 32% of correlations between ANME and non-ANME taxa are represented by OTUs assigned to three groups of sulfate-reducing bacteria: SEEP-SRB1g, SEEP-SRB1a, and SEEP-SRB2 (C). Stacked histograms of correlations between OTUs assigned to SEEP-SRB1g, SEEP-SRB1a, or SEEP-SRB2 and ANME OTUs, parsed by ANME subtype (D), highlights specific associations predicted between ANME-1 and either SEEP-SRB1a or SEEP-SRB2, ANME-2a and SEEP-SRB1a, ANME-2c and SEEP-SRB1a, and between ANME-2b and SEEP-SRB1g.

### mRNA imaging using HCR-FISH

Hybridization chain reaction FISH (HCR-FISH) is a powerful technique to amplify signal from FISH probes [65, 66]. The protocol used here was modified from Yamaguchi and coworkers [67]. *nifH* initiators, purchased from Molecular Technologies (Pasadena, CA, USA; probe identifier “nifH 3793/D933”) or designed in-house (Supp. Table 2) and ordered from Integrated DNA Technologies, were hybridized to fixed ANME-SRB consortia. Hairpins B1H1 and B1H2 with attached Alexa647 fluorophores (Molecular Technologies) were added separately to two 45 μL volumes of amplification buffer in PCR tubes and snap cooled by placement in a C1000 Touch Thermal Cycler (BioRad, Hercules, CA, USA) for 3 min at 95°C. After 30 min at room temperature, hairpins were mixed and placed in PCR tubes along with hybridized ANME-SRB consortia. Amplification was performed for 15 min at 35°C. Similar results were observed when the HCR-FISH v3.0 protocol established by Choi et al. [68] was used.

### Stable Isotope Probing and nanoSIMS

Methane seep sediments containing abundant ANME-2b and SEEP-SRB1g consortia (Supp. Fig. 4) were used in stable isotope probing (SIP) experiments to test for diazotrophic activity by SEEP-SRB1g. N sources were removed from the sediment slurry by washing with artificial seawater without an N source (see Supplemental Materials and Methods). Two anoxic incubations were pressurized with 2.8 bar CH_4_ with 1.2 mL ^15^N_2_ at 1 bar, approximately equivalent to 2% headspace in 20 mL CH_4_ at 2.8 bar (Supp. Table 3). Positive control incubations (*n* = 2) were amended with ^15^NH_4_Cl and were further pressurized with 2.8 bar CH_4_ and 1.2 mL natural-abundance N_2_ at 1 bar. Incubations were periodically checked for AOM activity via sulfide production using the Cline assay [69] and were chemically fixed for FISH-nanoSIMS analysis [70] after 9 months. Fixed ANME-SRB consortia were separated from the sediment matrix and concentrated following published protocols [5]. Samples were then embedded in Technovit H8100 (Kulzer GmbH, Germany) resin according to published protocols [5, 31] and thin sections (2 μm thickness) were prepared using an Ultracut E microtome (Reichert AG, Austria) which were mounted on Teflon/poly-L-lysine slides (Tekdon Inc., USA). FISH reactions were performed using Seep1g-1443 and ANME-2b-729 probes as described above, with the omission of 10% SDS to prevent detachment of section from slide (G. Chadwick, pers. comm.), and slides were imaged and mapped for subsequent nanoSIMS analysis using a Zeiss Elyra PS.1 platform. After removal of DAPI-Citifuor by washing following published protocols [70], slides were cut to fit into nanoSIMS sample holders and sputter-coated with 40 nm Au using a Cressington sputter coater. NanoSIMS was performed using a Cameca NanoSIMS 50L housed in Caltech’s Microanalysis Center, and data was analyzed using look@nanoSIMS [71].

## Results

### 16S rRNA correlation analysis predicts a specific association between ANME-2b and SEEP-SRB1g

Correlation analysis applied to 16S rRNA gene amplicon libraries has been frequently used to identify interactions between microorganisms based on the co-occurrence of their 16S rRNA sequences in different environments or conditions [72–75]. Here, we applied correlation analysis to 16S rRNA amplicon libraries prepared from Costa Rican methane seep sites (Supp. Table 1) to explore partnership specificity between ANME and associated SRB. QIIME processing of amplicon sequences prepared from 310 Costa Rican methane seep sediment and seep carbonate samples yielded 3,052 OTUs after filtering in R. A table of read abundances for these OTUs across the 310 samples was analyzed by SparCC to calculate correlation coefficients and significance for all possible 4,658,878 OTU pairs using 100 bootstraps (Fig. 1). Of these pairs, 9.7% (452,377) had pseudo-*p*-values < 0.01, indicating the coefficients for each of these correlations exceeded that calculated for that same OTU pair in any of the 100 bootstrapped datasets [41]. The taxonomic assignment of the constituent OTUs of correlations with pseudo-*p* < 0.01 were then inspected, where 18% (81,459) of correlations with pseudo-*p* < 0.01 describe those involving ANME. Of these, 32% occur between ANME and OTUs assigned to three main taxa: *Desulfococcus* sp., SEEP-SRB1a, and SEEP-SRB2 (Fig. 1). A complete list of significant correlations, their coefficient values, OTU identifiers, and accompanying taxonomy assignments can be found in Supplemental Table 4.

16S rRNA phylogenetic analysis revealed the SILVA-assigned “*Desulfococcus* sp.” OTUs comprise a sister clade to the SEEP-SRB1a that is distinct from cultured *Desulfococcus* sp. (e.g. *D. oleovorans* and *D. multivorans*, see below). We therefore reassigned the *Desulfococcus* OTUs to a new clade we termed SEEP-SRB1g following the naming scheme outlined for seep-associated SRB in Schreiber, et al. (e.g. SEEP-SRB1a through -SRB1f) [15]. Furthermore, statistically-significant correlations between OTUs of ANME and SRB taxa suggested that ANME-SRB partnerships in the Costa Rica seep samples could be classified into the following types: ANME-1 with SEEP-SRB1a or SEEP-SRB2, ANME-2a with SEEP-SRB1a, ANME-2b with SEEP-SRB1g, ANME-2c with SEEP-SRB1a or SEEP-SRB2, and ANME-3 with SEEP-SRB1a (Fig. 1). While physical association between different ANME lineages and Deltaproteobacterial clades SEEP-SRB1a and SEEP-SRB2 had been well-documented [5, 13, 15, 31, 34], members of the SEEP-SRB1g had not previously been identified as a potential syntrophic partner with methanotrophic ANME.

These associations were further examined by detailed network analysis in which the table of correlations with pseudo *p*-values < 0.01 was further filtered to contain only those correlations with coefficients (a measure of correlation strength) in the 99^th^ percentile of all significant correlations. A network diagram in which nodes represent OTUs and edges between nodes represent correlations was constructed with force-directed methods [76], where edge length varied in inverse proportion to correlation strength. A subregion of this network focused on ANME-associated OTUs is presented in Figure 2a. Cohesive blocks, subsets of the graph with greater connectivity to other nodes in the block than to nodes outside [77], were calculated and revealed 3 primary blocks of ANME and SRB OTUs. Visualization of these 3 blocks by a chord diagram [78] further highlighted the taxonomic identity of ANME-SRB OTU pairs in these blocks: ANME-1 or ANME-2c (one OTU with mean read count < 10) and SEEP-SRB2, ANME-2a or ANME-2c and SEEP-SRB1a, and ANME-2b or ANME-2a and SEEP-SRB1g (Fig. 2b). The predicted associations between ANME-2c and SEEP-SRB2 and between ANME-2a and SEEP-SRB1g were relatively more rare than the other associations; only one rare ANME-2c OTU (mean read count < 10) and four uncommon ANME-2a OTUs (mean read count < 100) were predicted between SEEP-SRB2 and SEEP-SRB1g, respectively. Inferred partnership specificity in two of the blocks has been previously corroborated by FISH studies, namely that exhibited by ANME-1 with SEEP-SRB2 [13, 34], ANME-2c with SEEP-SRB1a [5], and ANME-2a with SEEP-SRB1a [15]. The partnership between SEEP-SRB1g and ANME-2b, however, had no precedent, as the only previous FISH descriptions of ANME-2b had placed it with a partner Deltaproteobacterium with taxonomy not known beyond the phylum level [5, 31].

**Figure 2.**
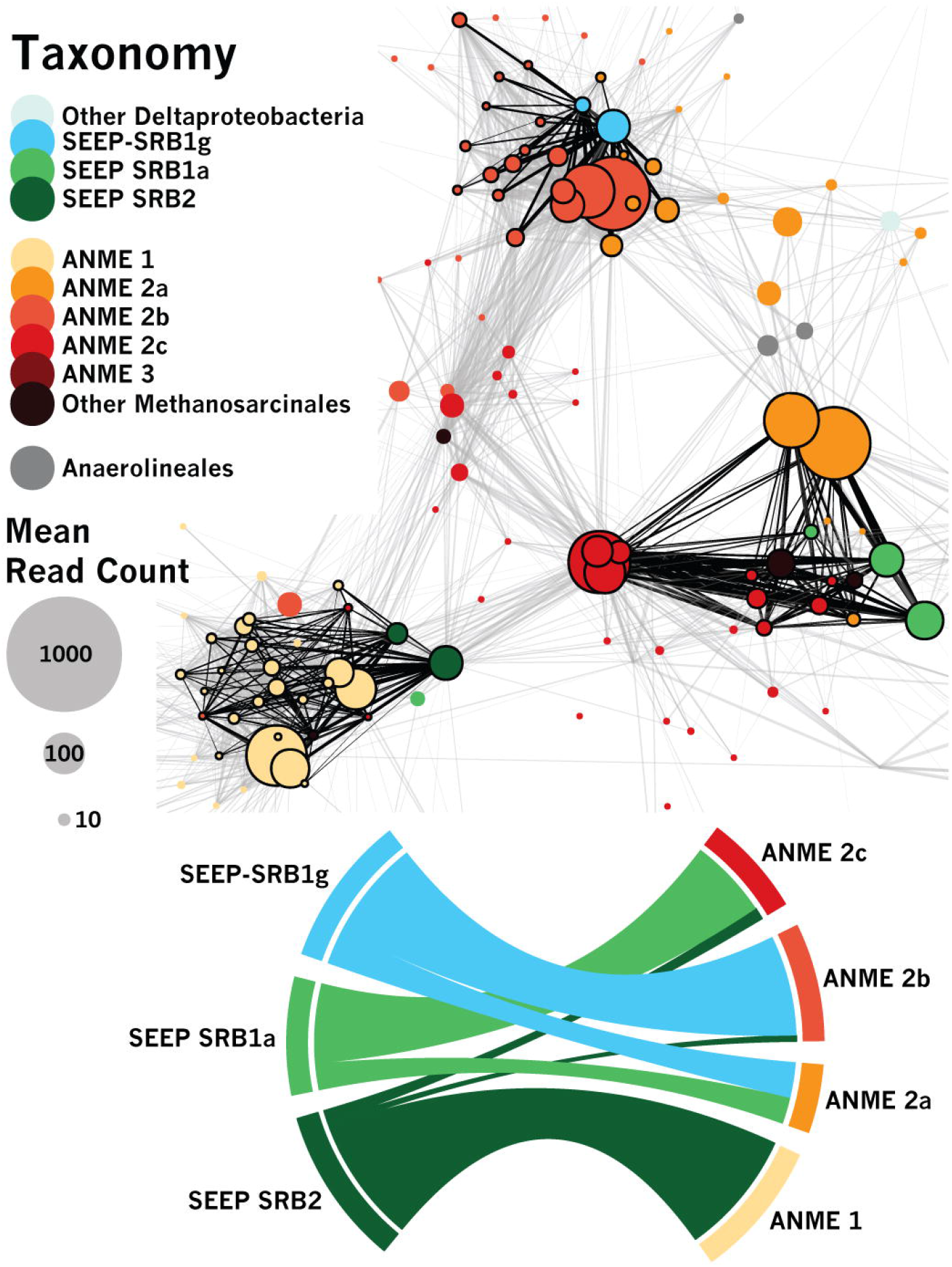
Network analysis of the subset of correlations between OTUs calculated by SparCC [41] that are both significant (*pseudo-p*-values < 0.01, 100 bootstraps) and strong (≥ 99^th^ percentile). Correlation strength is proportional to edge length and used to visualize the network (top panel) using force-directed methods [76]. Edges are black where they belong to a set of cohesive blocks of nodes [77] and gray otherwise. Chord diagram [78] visualizing ANME-SRB partnership specificity (bottom panel), with band thickness between SRB (left) and ANME (right) proportional to the number of edges between ANME and SRB OTUs within cohesive blocks. Network analysis supports (cf. Fig. 1) previously-undescribed association between ANME-2b and SEEP-SRB1g.

### Common patterns of association observed in network analysis and in single ANME-SRB consortia

To test if ANME-SRB partnership specificity observed in our iTag correlation analysis from seep samples (Figs. 1, 2) was consistent with data collected from individually-sorted ANME-SRB consortia after BONCAT-FACS [39], we constructed a phylogeny with full-length and amplicon 16S rRNA sequences from ANME-associated SRB including SEEP-SRB1g (Fig. 3; Supp. Fig 5). 16S rRNA iTag amplicon sequences from the network analysis (Fig. 2) and from BONCAT-FACS sorted consortia (Fig. 3; [39]) were then annotated by ANME subtype and identity of associated phylotypes. In the BONCAT-FACS dataset, 8 out of 11 (72%) of the consortia with ANME-2b OTUs had corresponding deltaproteobacterial OTUs that belonged to the SEEP-SRB1g clade (Fig. 3). Similarly, of the Deltaproteobacteria OTU sequences from the BONCAT-FACS sorted consortia affiliated with SEEP-SRB1g 89% (8/9) had ANME-2b as the archaeal partner (Fig. 3). Notably, we found that these SEEP-SRB1g sequences were also highly-similar to published full-length 16S rRNA clone library sequences (e.g. NCBI accession AF354159) from seep sediments where ANME-2b phylotypes were also recovered [21]. A *χ*^2^-test for independence was performed on 16S rRNA OTUs recovered from (39) to test the null hypothesis that the presence of a given SRB taxon in a FACS sort is independent of the type of ANME present in the sort. This test demonstrated that the SRB taxon found in a given sort was dependent on the ANME also present in the sort, *χ*^2^ = 30.6 (*d.f.* = 6, *n* = 30), *p* < 0.001. The pattern of association between ANME and SRB OTUs in individual BONCAT-FACS-sorted ANME-SRB consortia thus corroborated the inference from network analysis that ANME-2b and SEEP-SRB1g OTUs exhibit significant partnership specificity. On the basis of amplicon sequence associations found from the BONCAT-FACS sorting dataset as well as those displayed by correlation analysis of amplicons from Costa Rica methane seeps, we designed a set of independent experiments to test the hypothesis that ANME-2b form syntrophic partnerships with the previously-undescribed SEEP-SRB1g deltaproteobacteria.

**Figure 3.**
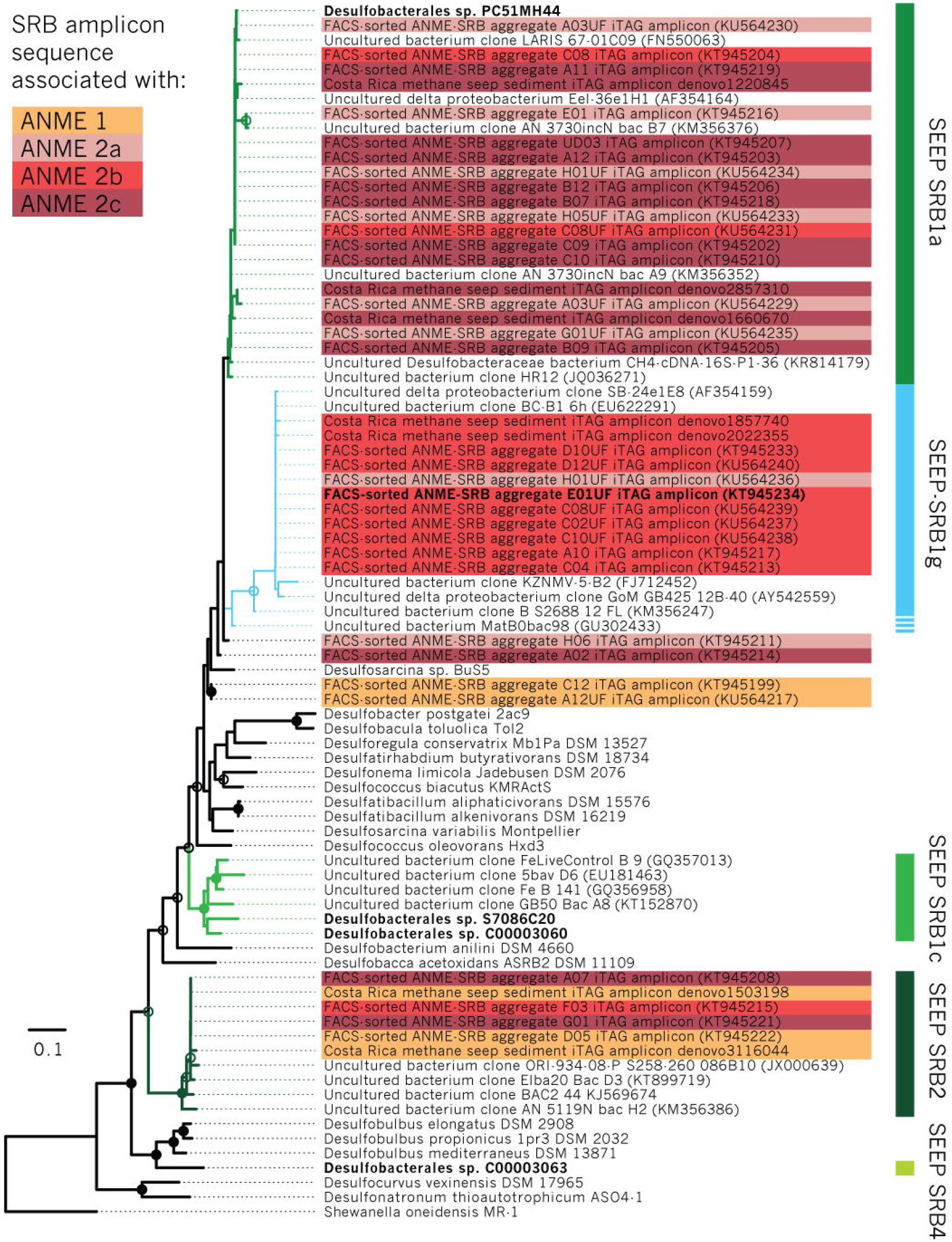
16S rRNA phylogenetic tree of methane seep Deltaproteobacteria and other lineages, including sequences from recovered metagenome-assembled genomes (MAGs) [14], iTag amplicons from BONCAT-FACS-sorted ANME-SRB consortia [39], iTag amplicon data from this study, and previously published clone library sequences. Maximum likelihood phylogeny was inferred using 100 bootstraps with >70% or 90% bootstrap support of internal nodes indicated with open or closed circles, respectively. Taxa associated with SRB amplicon iTag sequences were determined from data in Hatzenpichler, et al. 2016 [39] (BONCAT-FACS-sorted ANME-SRB consortia), and by network analysis of iTag amplicon data from methane seep samples (cf. Fig. 2). Taxa in bold represent 16S rRNA sequences from MAG bins acquired from methane seep sediments [14] or from BONCAT-FACS-sorted ANME-SRB consortia, including associated iTag amplicons [39]. The SEEP-SRB1a and −1g clades are operationally defined here by the extent of matches to the respective 16S rRNA FISH probes Seep1a-1441 and Seep1g-1443. Given the low bootstrap values for divergent sequences, the true extent of the SEEP-SRB1g clade is unclear, indicated by the dashed line (cf. Supp. Fig. 5).

### FISH experiments show SEEP-SRB1g in association with ANME-2b

Specific oligonucleotide probes were designed and tested for the SEEP-SRB1g clade (Supp. Fig. 1) and FISH experiments were used to validate the predicted ANME-2b—SEEP-SRB1g partnership. Simultaneous application of FISH probes targeting SEEP-SRB1a, the dominant deltaproteobacterial partner of ANME (Seep1a-1441 [15]), the newly designed SEEP-SRB1g probe (Seep1g-1443, this work), and a probe targeting ANME-2b (ANME-2b-729 [39]) demonstrated that ANME-2b predominantly form consortia with SEEP-SRB1g, appearing as large multicellular consortia in seep sediment samples from different localities at Costa Rica methane seep sites (see Supplemental Materials and Methods for site details) that also contain ANME-2a (Fig. 4b) and ANME-2c (Fig. 4f). ANME-2b was not observed in association with SEEP-SRB1a (Figs. 4a, 4e), and SEEP-SRB1g was not observed in association with ANME-2a (Fig. 4d) or ANME-2c (Fig. 4h) when FISH probes ANME-2a-828 or ANME-2c-760 [20] were substituted for ANME-2b-729 (*n* ≈ 100 consortia). Instead, SEEP-SRB1a was found in consortia with ANME-2a (Fig. 4c) and ANME-2c (Fig. 4g), consistent with previous reports [15].

**Figure 4.**
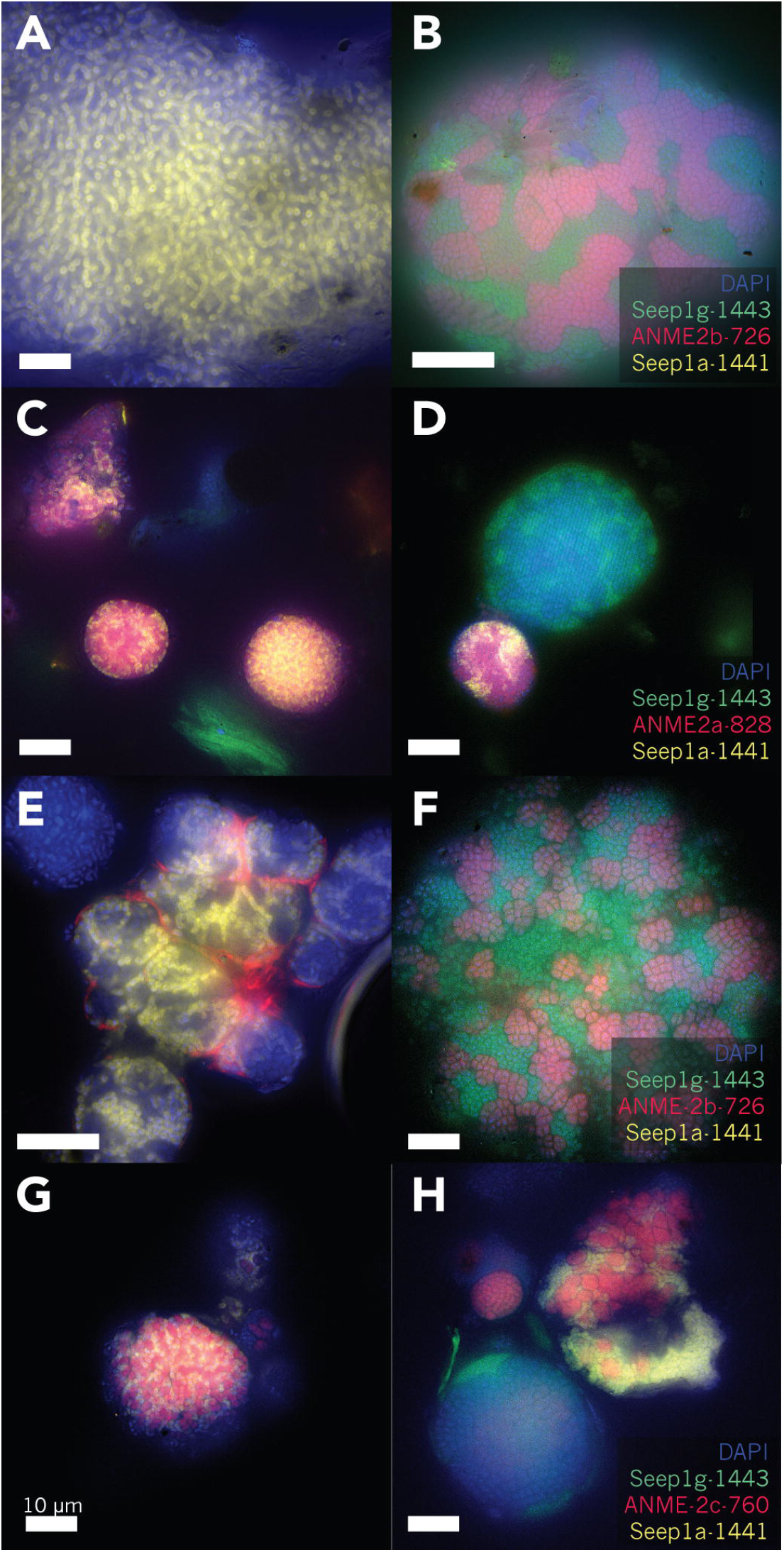
FISH data targeting AOM consortia in seep sediment samples using oligonucleotide probes targeting ANME-2b (ANME-2b-726) and ANME-2a (ANME-2b-828); (in red), a SEEP-SRB1a (Seep1a-1443) probe (in green) and a newly-designed probe (Seep1g-1443) targeting the SEEP-SRB1g clade (in yellow) demonstrating physical association between ANME-2b and SEEP-SRB1g. DAPI counterstain is shown in blue. Examples of ANME-2b—SEEP-SRB1g (B, F) and ANME-2a/ANME-2c–SEEP-SRB1a (C, D, G, H) partnership specificity. Seep sediments harboring ANME-2a and ANME-2b (A-D) host AOM consortia that are composed primarily of either ANME-2a–SEEP-SRB1a or ANME-2b–SEEP-SRB1g (B, C, D). The absence of hybridization with the ANME-2b probe in AOM consortia positively hybridized by the SEEP-SRB1a probe (A) and absence of ANME-2a probe hybridization in SEEP-SRB1g-containing consortia (D) further supports distinct ANME-2—SRB pairings for ANME-2a and ANME-2b. Similarly, FISH analysis of AOM consortia from sediments rich in ANME-2c and ANME-2b (E-H) were composed almost entirely of ANME-2b–SEEP-SRB1g or ANME-2c–SEEP-SRB1a partnerships (F, G, H); AOM consortia positively hybridized with the SEEP-SRB1g or SEEP-SRB1a probes were not observed to hybridize with probes targeting ANME-2c (H) or ANME-2b (E), respectively. In all panels, the scale bar is 10 μm.

### Genomic potential for N_2_ fixation in sulfate-reducing SEEP-SRB1g deltaproteobacteria

Given the importance of diazotrophy in the functioning of ANME-SRB syntrophy, we screened metagenome-assembled genome bins (MAGs) of SEEP-SRB1g for the presence of the nitrogenase operon. A genome tree constructed from previously published MAGs from Hydrate Ridge and Santa Monica Basin [14, 39] revealed that two closely related MAGs (Desulfobacterales sp. C00003104, and C00003106) originally classified as belonging to the Seep-SRB1c clade [14] possessed the nitrogenase operon (Fig. 5). These MAGs did not possess 16S rRNA genes, precluding 16S rRNA-based taxonomic identification. A more detailed look at these reconstructed genomes revealed that the nitrogenase along with a suite of other genes were unique to this subclade and missing in other SEEP-SRB1c MAGs [14], suggesting they may represent a distinct lineage. In effort to connect these nitrogenase containing SRB MAG’s with representative 16S rRNA sequences, we examined mini-metagenome data from individual BONCAT-FACS sorted ANME-SRB consortia which each contained 16S rRNA gene sequences for the ANME and bacterial partner [39]. A genome tree containing deltaproteobacterial MAGs from Skennerton, et al. [14] and reconstructed deltaproteobacterial genomes from the BONCAT-FACS sorts [39] revealed one SRB genome from a FACS-sorted consortium (Desulfobacterales sp. CONS3730E01UFb1, IMG Genome ID 3300009064) was closely related to the two putative Seep-SRB1c MAGs containing the nitrogenase operon (Fig. 5). The 16S rRNA amplicon sequence (NCBI accession KT945234) associated with the Desulfobacterales sp. CONS3730E01UFb1 genome was used to construct a 16S rRNA phylogeny and confirmed to cluster within the SEEP-SRB1g clade, providing a link between the 16S rRNA and associated nitrogenase sequences in this lineage (Fig. 3). Given that Desulfobacterales sp. CONS3730E01UFb1, C00003104, and C00003106 genomes appeared highly similar on the genome tree (Fig. 5), we reassigned the previously published Desulfobacterales sp. C00003104 and C00003106 MAGs to the SEEP-SRB1g. Notably, the other 16S rRNA amplicon sequence sampled from the sorted consortium CONS3730E01UF (NCBI accession KT945229) was assigned to ANME-2b [39].

**Figure 5.**
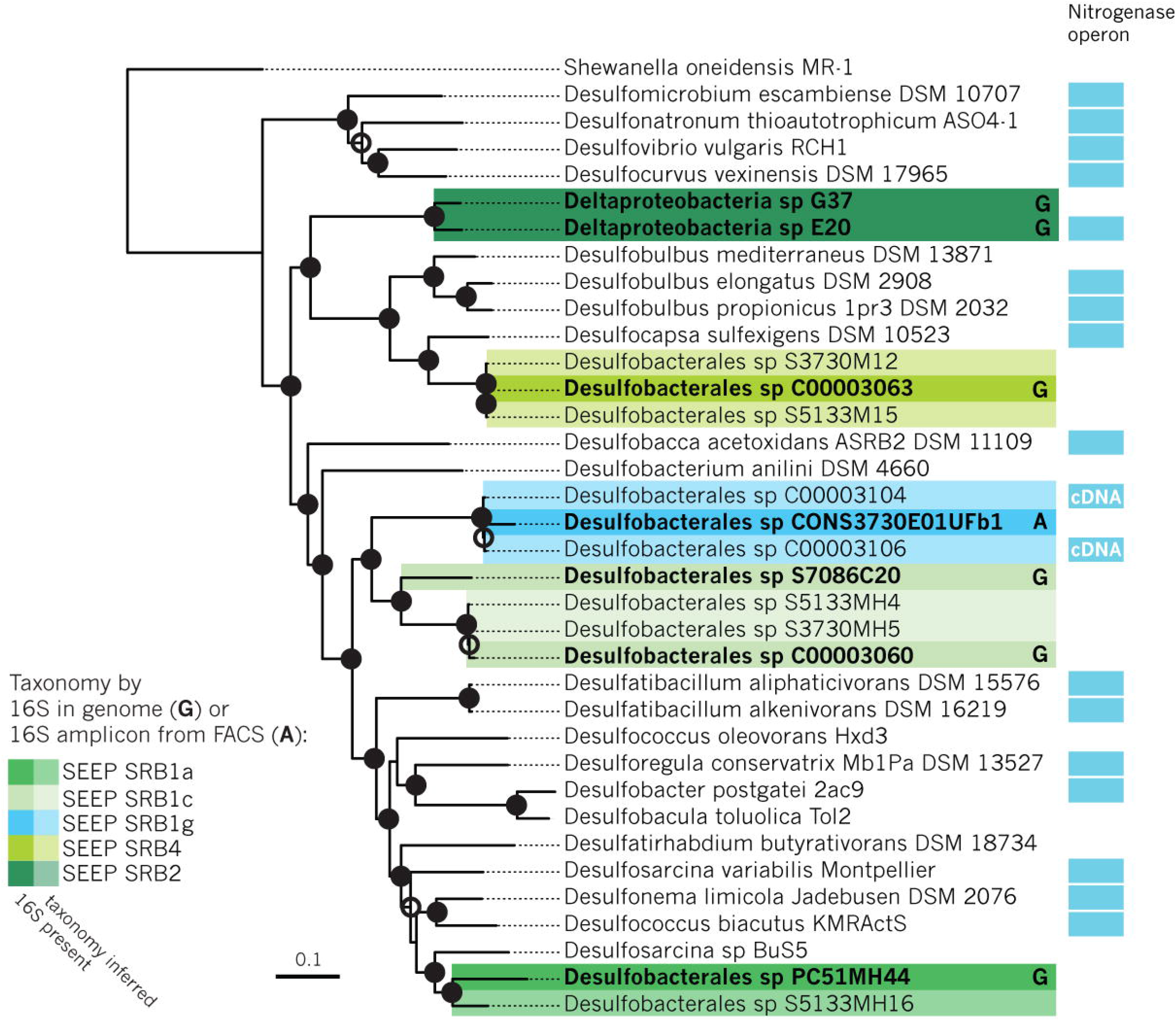
Genome tree of ANME-associated Deltaproteobacteria and related organisms inferred from maximum likelihood methods. Bootstrap support for internal nodes was determined using 100 bootstraps and depicted on the tree as open (>70% bootstrap support) or closed (>90%) circles. Genome bins containing a 16S rRNA gene or an associated 16S iTag amplicon sequence are highlighted in bold and with a color corresponding to 16S taxonomy assignment. Inferred taxonomy of genome bins closely related to bins containing 16S rRNA sequences are highlighted in a lighter shade. Genome bins containing the nitrogenase operon are annotated with a blue bar. *nifH* sequences found to be expressed in methane seep sediments in cDNA clone libraries [8] are annotated by “cDNA”. As noted in the text, a search of unpublished SEEP-SRB1a MAGs revealed the presence of highly-expressed [8] *nif*H sequences in several unpublished bins (Supp. Fig. 6).

As noted above, these SEEP-SRB1g MAGs were remarkable for the presence of the *nifHDK* operon involved nitrogen fixation, which had previously not been an area of focus in previous analyses of ANME-associated SRB genomes (Fig. 5). A re-analysis of published *nifH* cDNA sequences from methane seep sediments revealed sequences that were nearly identical to the SEEP-SRB1g *nifH* (NCBI accession KR020451-KR020457, [8]) suggesting active transcription of SEEP-SRB1g *nifH* under *in situ* conditions (Fig. 6). An analysis of published methane seep metaproteomic data [14] also indicated active translation of nitrogenase by SEEP-SRB1g, corroborating evidence from cDNA libraries [8]. Additionally, other *nifH* cDNA sequences in this study were found to be identical to nitrogenase sequences occurring in 18 SEEP-SRB1a unpublished metagenome bins (Supp. Fig. 6) demonstrating that at least some of the syntrophic SEEP-SRB1a SRB partners also possess and actively express *nifH*.

**Figure 6.**
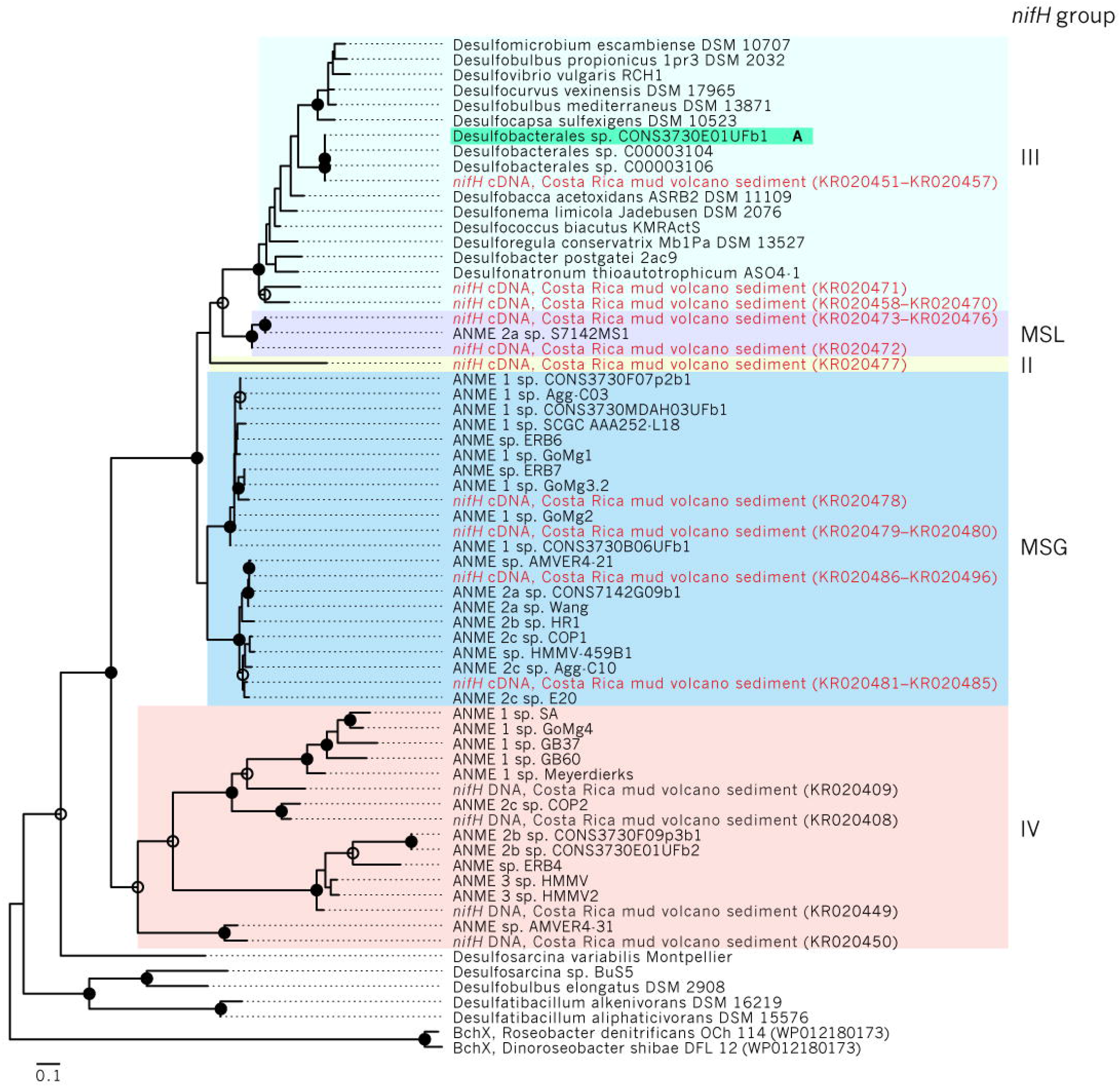
Phylogeny of *nifH* sequences extracted from *nifH* cDNA (red text) and DNA clone libraries [8], from genome bins acquired from methane seep sediments [14], and from other Deltaproteobacteria genomes using a tblastn search with chlorophyllide reductase BchX (WP011566468) as a query. This BchX sequence along with another BchX (WP012180173) were used as an outgroup to root the tree. Phylogeny was inferred by maximum likelihood methods using 100 bootstraps; bootstrap support of internal nodes is illustrated as open or closed circles, indicating >70% or >90% bootstrap support, respectively. *nifH* recovered from the BONCAT-FACS-sorted genome CONS3730E01UFb1, a bin with an accompanying 16S rRNA amplicon sequence placing it within the SEEP-SRB1g, is highlighted in teal. *nifH* groups (sensu Raymond et al. [95]) were assigned by comparison with Dekas, et al. 2016, and are annotated either by group number or abbreviated as follows: MSL, Methanosarcina-like; MSG, Methane Seep Group.

### Single-cell nifH expression visualized by HCR-FISH and ^15^N_2_ FISH-nanoSIMS experiments confirm involvement of SEEP-SRB1g in N_2_-fixation

The dominant role of ANME-2 in nitrogen fixation reported by previous studies [8–10] motivated our examination of whether the sulfate-reducing SEEP-SRB1g partners of ANME-2b were also involved in diazotrophy, either in concert with the ANME-2b partner, or perhaps as the sole diazotroph in this AOM partnership. Using the *nifH* sequences from SEEP-SRB1g, we designed a specific mRNA-targeted probe set to use in whole-cell hybridization chain reaction FISH (HCR-FISH) assays (Supp. Table 2). HCR-FISH allows for signal amplification and improved signal-to-noise ratio compared to FISH, and has been used in single cell mRNA expression studies in select microbial studies [79–81]. Prior to this study, however, HCR-FISH had not been applied to visualize gene expression in ANME-SRB consortia from methane seep sediments. In the context of experiments with sediment-dwelling ANME-SRB consortia, HCR-FISH provided adequate amplification of the signal to detect expressed mRNA above the inherent background autofluorescence in sediments. Using our HCR-FISH probes targeting SEEP-SRB1g *nifH* mRNA together with the standard 16S rRNA targeted oligonucleotide FISH probes Seep1g-1443 (targeting SEEP-SRB1g) and ANME-2b-729 (targeting ANME-2b), we successfully imaged *nifH* mRNA transcripts by SEEP-SRB1g cells in ANME-2b—SEEP-SRB1g consortia in a sediment AOM microcosm experiment (Fig. 7). Positive HCR-FISH *nifH* hybridization in this sample was observed to be exclusively associated with the SEEP-SRB1g bacterial partner in ANME-2b consortia (*n* = 5), and not observed in ANME-2b stained cells nor in ANME-2a or −2c consortia, supporting the specificity of this assay. Negative control experiments for the HCR-FISH reaction were also performed in which SEEP-SRB1g *nifH* initiator probes were added to the assay, but the fluorescent amplifier hairpins were absent. In this case, there is no fluorescent signal in either the bacteria or archaeal partners in ANME-2b aggregates confirming that there is no native autofluorescence in Seep-SRB1g that could be responsible for the signal observed in the HCR-FISH experiments (Supp. Fig. 7f-j). We performed another control without the initiator probes that bind the mRNA but with the addition of the fluorescent amplifier hairpins. As in the other negative control, we observed limited non-specific binding of the hairpins that were easy to differentiate from the positively-hybridized SEEP-SRB1g (Supp. Fig. 7a-e). Occasionally, highly localized and small spots of hairpins were observed (Supp. Fig 7d) but these dots were mostly localized outside of aggregates and did not align with either bacteria or archaea in consortia (e.g. Figure 7d). Confirmation of *nifH* expression using HCR-FISH corroborated evidence from cDNA libraries (Fig. 6) that SEEP-SRB1g actively express *nifH*, providing support for diazotrophy in the sulfate-reducing partner in ANME-2b—SEEP-SRB1g consortia.

**Figure 7.**
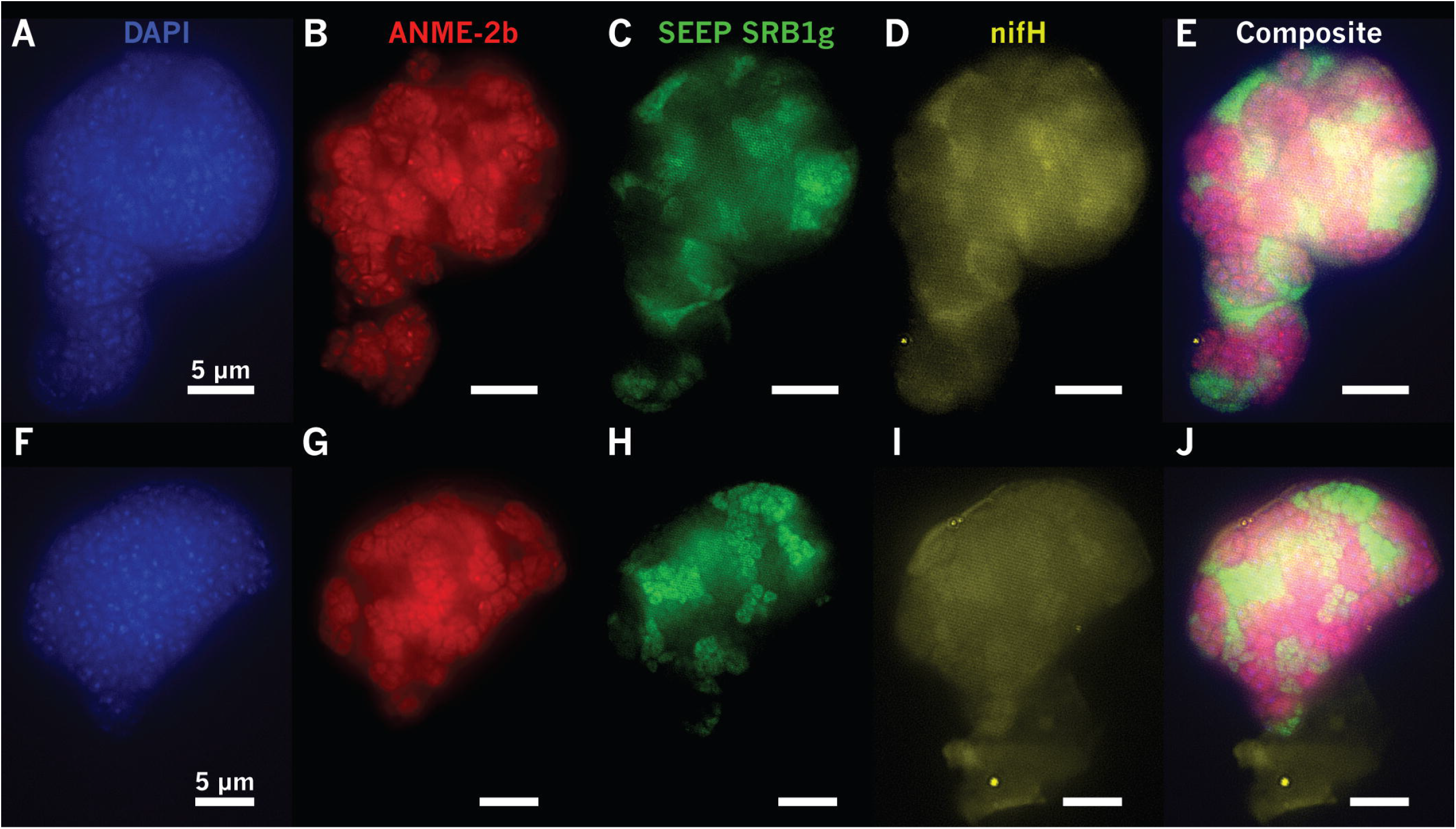
HCR-FISH assays show *in situ* expression of *nifH* in SEEP-SRB1g in association with ANME-2b in methane seep sediment incubations. ANME-2b (B, G) and SEEP-SRB1g (C, H) cells labeled with FISH probes ANME-2b-729 (in red, [39]) and newly-designed Seep1g-1443 (in green) with DAPI as the DNA counterstain (A,F). HCR-FISH targeting SEEP-SRB1g *nifH* mRNA (in yellow; Supp. Table 2) demonstrated active expression of *nifH* transcripts localized to SEEP-SRB1g cells (D, I), supporting the hypothesis of diazotrophy by partner SRB. Scale bars in all panels are 5 μm.

To confirm active diazotrophy by ANME-2b-associated SEEP-SRB1g, we prepared stable isotope probing incubations of methane seep sediments recovered from a Costa Rica methane seep. These nitrogen-poor sediment incubations were amended with unlabeled methane and ^15^N_2_ and maintained in the laboratory at 10°C under conditions supporting active sulfate-coupled AOM (see Supplemental Materials and Methods). Sediments with abundant ANME-SRB consortia were sampled after 9 months of incubation and separated consortia were analyzed by nanoSIMS to measure single cell ^15^N enrichment associated with diazotrophy within ANME-2b—SEEP-SRB1g consortia. Representative ANME-2b—SEEP-SRB1g consortia (*n* = 4) were analyzed by FISH-nanoSIMS and shown to be significantly enriched in ^15^N relative to natural abundance values (0.36%; Fig. 8). Among the consortia analyzed, the ^15^N fractional abundance in ANME-2b cells were often higher than that measured in SEEP-SRB1g, with ANME-2b cells on the exterior of an exceptionally large consortium (Fig. 8b-c) featuring ^15^N fractional abundance of 1.73% ± 0.14 (number of ROIs, *n* = 72), significantly enriched relative to that measured in SEEP-SRB1g cells in the exterior, 0.77% ± 0.09 (*n* = 58). This indicated that ANME-2b were often the primary diazotroph in consortia, consistent with previous reports from ANME-2–DSS consortia [8–11]. Notably, however, in one ANME-2b—SEEP-SRB1g consortium, the SEEP-SRB1g cells were more enriched in ^15^N relative to the associated ANME-2b cells, with ANME-2b cells containing 1.34% ± 0.13 ^15^N (*n* = 82) and SEEP-SRB1g containing 3.02% ± 0.20 ^15^N (*n* = 22, Fig. 8i), suggesting that under certain circumstances the sulfate-reducing partner may serve as the primary diazotroph. This pattern suggests diazotrophic flexibility in ANME-2b—SEEP-SRB1g consortia in which one partner–ANME-2b or SEEP-SRB1g–can serve as the primary diazotroph in the consortium. Additionally, a gradient in ^15^N enrichment in a the large ANME-2b consortium was observed in which clusters of ANME-2b cells associated with the interior of the consortia were significantly more enriched in ^15^N relative to ANME-2b clusters near the aggregate exterior, with ^15^N fractional abundances for ANME-2b cells in the exterior of 1.73% ± 0.14 (*n* = 72), significantly higher than those measured for ANME-2b cells in the interior, 2.64% ± 0.14 (*n* = 116). Notably, no equivalent gradient was observed in the SEEP-SRB1g partner, with SEEP-SRB1g cells in the exterior displaying ^15^N fractional abundances of 0.77% ± 0.09 (*n* = 58) compared with those measured on the interior, 0.78% ± 0.09 (*n* = 62).

**Figure 8.**
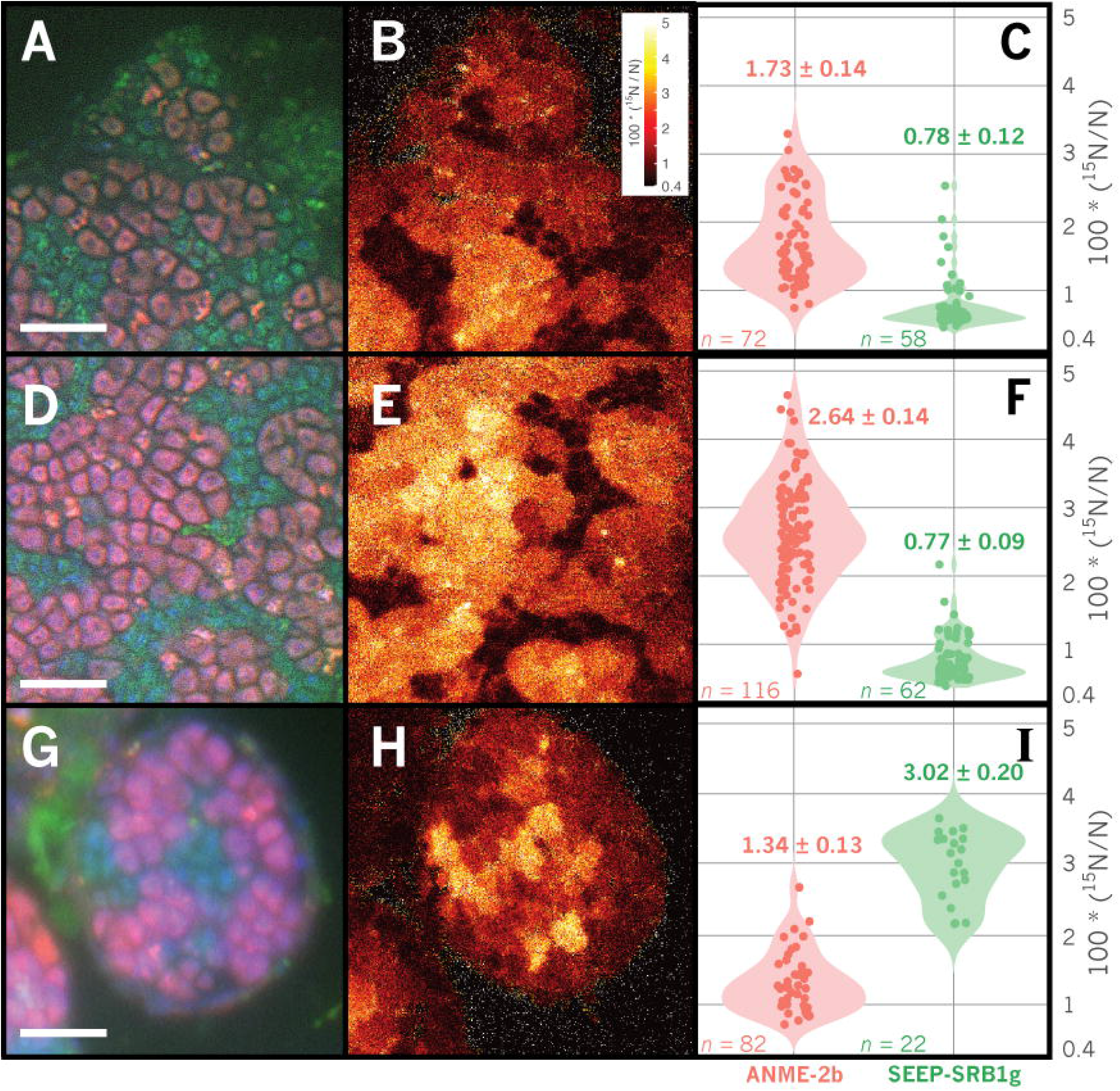
Correlated FISH-nanoSIMS imaging of representative ANME-2b–SEEP-SRB1g consortia demonstrating active diazotrophy by ANME-2b (B, E) and SEEP-SRB1g (H) cells through ^15^N incorporation from ^15^N_2_. FISH images of ANME-2b (pink) and SEEP-SRB1g (green) are shown in panels A, D, G and corresponding nanoSIMS ^15^N atom percent values are shown in panels B, E, and H. Scale bar is 5 μm in panels A, D, G; raster size in panels B, E, and H is 20 μm^2^. Violin plots (C, F, I) of ^15^N fractional abundance for each type of ROI, representing single ANME-2b or SEEP-SRB1g cells. The number of ROIs measured is indicated by *n* in each panel. Diazotrophic activity in ANME-2b cells appears to be correlated with spatial structure, evidenced by increasing ^15^N enrichment in cells located within consortia interiors (E, F). SEEP-SRB1g cells are also observed to incorporate ^15^N from ^15^N_2_, and appear to be the dominant diazotroph in the consortium shown in panels G, H, and I, with cellular ^15^N enrichment in SEEP-SRB1g cells greater than that of the paired ANME-2b partner. Abscissa minima set to natural abundance of ^15^N (0.36%).

## Discussion

The symbiotic relationship between ANME and associated SRB, originally described by Hinrichs [17], Boetius [4], and Orphan [21], has been the focus of extensive study using FISH [5, 7, 13, 15, 25, 26, 29, 34, 35], magneto-FISH [29, 37, 38], and BONCAT-FACS [39], techniques that have provided insight into the diversity of partnerships between ANME and SRB. While these fluorescence-based approaches offer direct confirmation of physical association between taxa and are thus useful for characterizing partnership specificity, they are often constrained by sample size and are comparatively lower-throughput than sequencing-based approaches. Next-generation Illumina iTag sequencing of 16S rRNA amplicon sequences offers advantages in terms of throughput and is rapidly becoming a standard approach in molecular microbial ecology studies. Correlation analysis performed on these large iTag datasets can be an effective hypothesis-generating tool for identifying microbial interactions and symbioses in the environment [75], but most studies employing this approach stop short of validating predictions. As correlation analysis of iTag datasets is known to be sensitive to false positives due to the compositional nature of 16S rRNA amplicon libraries [41, 42, 82], specific correlations predicted between taxa should be corroborated when possible by independent approaches.

In this study, we used correlation analysis of 16S rRNA iTag data from 310 methane seep sediment and carbonate samples on the Costa Rican Margin to identify well-supported (pseudo-*p*-values < 0.01) positive correlations between specific OTUs commonly observed in seep ecosystems. Our analysis identified strong correlations between syntrophic partners previously described in the literature, such as that between members of the SEEP-SRB1a and ANME-2a/ANME-2c clades and between ANME-1 and SEEP-SRB2 [5, 7, 13, 15, 25, 26, 29, 34, 35], and uncovered previously unrecognized relationships between members of the ANME-2b clade and OTUs affiliated with an uncultured Desulfobacterales lineage, SEEP-SRB1g (Figs. 1-3). We then validated the specificity of the ANME-2b and SEEP-SRB1g association by FISH (Fig. 4).

The specificity of the association between ANME-2b and SEEP-SRB1g appeared to extend beyond Costa Rica methane seeps and is likely a widespread phenomenon, as this association was also recovered from BONCAT-FACS datasets originating from methane seep sites off of Oregon, USA (Hydrate Ridge) and from the Santa Monica Basin, California, USA. Our observations of ANME-2b—SEEP-SRB1g partnership specificity in numerous samples is consistent with published observations of other ANME-SRB partnerships, where consortia composed of specific ANME and SRB clades have been observed in seep ecosystems worldwide [15]. Notably, the syntrophic relationship between ANME-2b and SEEP-SRB1g appears to be highly specific (Fig. 2), as FISH observations from sediment samples from multiple Costa Rica methane seep sites (Supp. Table 1) did not show ANME-2b in consortia with other bacteria besides the SEEP-SRB1g (Fig. 4). This is in contrast with SEEP-SRB1a which, in these same experiments, was found to form associations with both ANME-2a and ANME-2c, indicative of this SRB syntroph having a broader capacity for establishing associations with methanotrophic ANME. Members of the diverse ANME-2c lineage also appeared to display partnership promiscuity in our network analysis, with positive correlations observed between ANME-2c OTUs and both SEEP-SRB1a and SEEP-SRB2 OTUs (Fig. 2). This predicted partnership flexibility in the network analysis was again corroborated by FISH observations of ANME-2c—SEEP-SRB1a consortia (Fig. 4) and prior reports of ANME-2c in association with SEEP-SRB2 from Guaymas Basin sediments [13]. These collective data suggest that partnership specificity varies among different clades of ANME and SRB, which may be the result of physiological differences and/or molecular compatibility, signal exchange, and recognition among distinct ANME and SRB that shape the degree of specificity between particular ANME and SRB partners, as has been observed in other symbiotic associations [83–85]. The degree of promiscuity or specificity for a given syntrophic partner may be influenced by the co-evolutionary history of each partnership, with some ANME or SRB physiologies requiring obligate association with specific partners. A more detailed examination of the genomes of ANME-2b and SEEP-SRB1g alongside targeted ecophysiological studies may provide clues to the underlying mechanism(s) driving specificity within this ANME-SRB consortia. Comparative investigations with ANME-2a and −2c subgroups may similarly uncover strategies enabling broader partner association, perhaps with preference for a SRB partner shaped by environmental variables rather than through pre-existing co-evolutionary relationships.

An initial genomic screening of SEEP-SRB1g offered some insight into the distinct metabolic capabilities of the SRB partner which may contribute to the association with ANME-2b. The observation of a complete nitrogenase operon in 3 different SEEP-SRB1g genome bins suggested the potential for nitrogen fixation, a phenotype not previously described for ANME-associated SRB (Fig. 5). While previous work on nitrogen utilization by ANME-SRB consortia has focused on diazotrophy performed by ANME-2 [8–10], environmental surveys of seep sediments have noted active expression of nitrogenase typically associated with Deltaproteobacteria [8, 86]. In these studies, the specific microorganisms associated with the expressed nitrogenase in methane seep sediments were not identified. Prior to our findings presented here, diazotrophy by ANME-associated SRB had not been demonstrated. A phylogenetic comparison of the *nifH* sequences associated with SEEP-SRB1g with sequences of the expressed deltaproteobacterial-affiliated (i.e. Group III) *nifH* transcripts reported in Dekas, et al. [8] revealed a high degree of sequence similarity, with SEEP-SRB1g related *nifH* among the most highly expressed (Figs. 5-6). Explicit tests for nitrogenase expression using HCR-FISH and active diazotrophy using stable isotope probing and FISH-nanoSIMS confirmed the involvement of SEEP-SRB1g in nitrogen fixation. Of the 4 ANME-2b—SEEP-SRB1g consortia analyzed by FISH-nanoSIMS, one had significantly more ^15^N enrichment in the SEEP-SRB1g partner relative to the ANME-2b, while the other 3 displayed higher cellular ^15^N enrichment in the ANME-2b partner (Fig. 8). This pattern supported our inference of diazotrophic flexibility within ANME-2b—SEEP-SRB1g consortia in which either the ANME or the SRB partner can serve as the primary diazotroph in the consortium. Additionally, our detection of nitrogenase operons in the reconstructed genomes of the dominant syntrophic SRB partner, SEEP-SRB1a (Supp. Fig. 6), suggests the potential for nitrogen fixation may extend to other bacterial partners as well and merits further investigation. Re-examination of nitrogen fixation in these partnerships with new FISH probes and nanoSIMS at single-cell resolution will further illuminate the full diversity of diazotrophic activity among ANME-SRB consortia and the associated environmental/ physiological controls.

The factors responsible for determining which partner becomes the primary diazotroph in ANME-2b—SEEP-SRB1g consortia requires targeted study, but our preliminary data suggest this may be influenced in part by the relative position of ANME-2b or SEEP-SRB1g cells, particularly within large (>50 μm) ANME-2b—SEEP-SRB1g consortia. Previous studies of nitrogen fixation in ANME-SRB consortia found no correlation between consortia size and diazotrophic activity in consortia with diameters < 10 μm [10], but larger consortia such as those presented here have not been examined at single-cell resolution. Additionally, consortia with the morphology observed here, in which ANME-2b cells form multiple sarcinal clusters surrounded by SEEP-SRB1g (Figs. 4b, 8), have not been the specific focus of nanoSIMS analysis but appear to be the common morphotype among ANME-2b—SEEP-SRB1g consortia [31]. The frequency with which this morphotype is observed in ANME-2b—SEEP-SRB1g consortia may be related to the underlying physiology of this specific partnership. NanoSIMS analysis of a particularly large ANME-2b—SEEP-SRB1g consortium (~200 μm) with this characteristic morphology (Fig. 8a-f) revealed a gradient in diazotrophic activity in which ANME-2b cells located in the interior of the consortium incorporated far more ^15^N from ^15^N_2_ than ANME-2b cells near the exterior. This pattern may be related to variations in nitrogen supply from the external environment, as similar patterns of nutrient depletion with increasing depth into microbial aggregates have been predicted in modeling studies of nitrate uptake in *Trichodesmium* sp. [87] and directly observed by SIMS in stable isotope probing studies of carbon fixation in biofilm-forming filamentous cyanobacteria [88]. In these examples, modeling and experimental results document declining nitrate or bicarbonate ion availability inwards toward the center of the aggregates resulting from nitrate or bicarbonate consumption. An analogous process may occur in large ANME-2b—SEEP-SRB1g consortia, where cells situated closer to the exterior of the consortium assimilate environmental NH_4_^+^, increasing nitrogen limitation for cells within the consortium core. Interestingly, the single consortium in which the SEEP-SRB1g partner was the inferred primary diazotroph featured SEEP-SRB1g cells in the core of this consortium with ANME-2b cells toward the exterior (Fig. 8). The current nanoSIMS dataset is small and determining the biotic and environmental factors that influence which partner serves as the primary diazotroph in ANME-2b—SEEP-SRB1g consortia necessitates further study, but a reasonable hypothesis is that the proximity of cells in a given ANME-2b—SEEP-SRB1g consortium relative to the consortium exterior (and NH_4_^+^ availability in the surrounding pore fluid) influences the spatial patterns of diazotrophic activity by both ANME and SRB in large consortia. The concentration of ammonium in seep porefluids can be highly variable over relatively small spatial scales (e.g. between 50 - 300 μM [80]), and rates of diazotrophy estimated from laboratory incubations of methane seep sediment samples indicate different threshold concentrations of NH_4_^+^_(aq)_ above which diazotrophy ceases, as low as 25 μM [89] to 100-1000 μM [90–92]. In the large consortia observed here, this threshold [NH_4_^+^_(aq)_] may be crossed within the consortium as NH_4_^+^ is assimilated by cells at the consortium exterior, inducing nitrogen limitation and diazotrophy by ANME or SRB near the consortium core. Given the importance of diazotrophy in ANME-SRB consortia for nitrogen cycling at methane seep communities [10, 89], future work should test this hypothesis with SIP incubations with ^15^N_2_ under variable [NH_4_^+^_(aq)_].

The observed variation in diazotrophic activity in ANME-2b or SEEP-SRB1g cells may also be the result of phenotypic heterogeneity [93] within the multicellular ANME-2b—SEEP-SRB1g consortia, in which expression of the nitrogenase operon that ANME-2b and SEEP-SRB1g partners both possess is an emergent behavior resulting from the spatial organization of ANME-2b and SEEP-SRB1g cells within the consortium. On the basis of nanoSIMS observations of heterogeneous diazotrophy in clonal *Klebsiella oxytoca* cultures, phenotypic heterogeneity was inferred to confer selective advantage on microbial communities by enabling rapid response to environmental fluctuations [94]. Similar heterogeneity in *nif* expression by ANME-2b or SEEP-SRB1g cells may provide partners with resilience against changes in environmental nitrogen supply. Corroborating these observations in diverse ANME-SRB consortia and direct coupling of single-cell mRNA expression with nanoSIMS-acquired ^15^N enrichment would further inform the degree to which relative arrangement of the partners and spatial structure within a consortium plays a significant role in determining the mode of nutrient or electron transfer between partners.

## Conclusions

Here, we present an effective approach to detect novel pairings of microbial symbionts by coupling correlation analysis of 16S rRNA amplicon libraries with FISH and BONCAT-FACS experiments, going beyond amplicon sequencing-based hypothesis generation to experimental validation of hypothesized partnerships using microscopy and single-cell sorting techniques. Correlation analysis performed on a 16S amplicon survey of methane seep sediments near Costa Rica uncovered a novel and highly-specific ANME-SRB partnership between ANME-2b and SEEP-SRB1g. This partnership specificity was then validated by FISH, and further corroborated by 16S rRNA amplicon sequences from BONCAT-FACS-sorted single ANME-SRB consortia from methane seep sediments near Costa Rica, Hydrate Ridge, and Santa Monica Basin in California. Preliminary genomic screening of representative genomes from SEEP-SRB1g uncovered potential for nitrogen fixation in these genomes. Examination of published *nifH* cDNA clone libraries [8] and transcriptomic data [14] prepared from methane seep sediments demonstrated that SEEP-SRB1g actively expresses *nifH in vivo*. Co-localization of signal for *nifH* mRNA and SEEP-SRB1g 16S rRNA by HCR-FISH further corroborated active transcription of *nifH* by SEEP-SRB1g. FISH-nanoSIMS analysis of ANME-2b—SEEP-SRB1g consortia grown with ^15^N_2_ headspace documented ^15^N incorporation in SEEP-SRB1g cells, suggesting that SEEP-SRB1g may fix nitrogen as well. Future work should focus on examining unique aspects of each ANME-SRB partnership to improve our understanding of the diversity of anaerobic methane oxidation symbioses endowed by evolution.

## Acknowledgements

The authors would like to acknowledge the ROC-HITS science party, R/V *Atlantis* crew and HOV *Alvin* pilots from AT37-13. We would like to thank G. Chadwick for helpful comments and H. Yu for assistance with sediment incubations. We additionally acknowledge Y. Guan for his assistance with the nanoSIMS analysis, R. Hatzenpichler for early BONCAT-FACS experiments, and M. Aoki (National Institute of Technology, Wakayama College, Japan) for design of the FISH probe ANME-2a-828. We would further like to thank M. Schwarzkopf and Molecular Technologies for designing a set of HCR-FISH probes for *nifH* mRNA. Funding for this work was provided by the US Department of Energy’s Office of Science (DE-SC0020373), the National Science Foundation BIO-OCE grant (#1634002) and the NSF-supported Center for Dark Energy Biosphere Investigations, and a Gordon and Betty Moore Foundation Marine Microbiology Investigator grant (#3780); (all to V.J.O.). A portion of this research was performed under the Facilities Integrating Collaborations for User Science (FICUS) initiative and used resources at the DOE Joint Genome Institute and the Environmental Molecular Sciences Laboratory, which are DOE Office of Science User Facilities. Both facilities are sponsored by the Office of Biological and Environmental Research and operated under Contract Nos. DE-AC02-05CH11231 (JGI) and DE-AC05-76RL01830 (EMSL). K.S.M. was supported in part by a National Science Foundation Graduate Research Fellowship and a Schlanger Ocean Drilling Fellowship. V.J.O. is a CIFAR Fellow in the Earth 4D: Subsurface Science and Exploration Program.

---

Supplemental Table 1. Samples of methane seep sediment used in this study to produce 16S rRNA amplicon libraries.

Supplemental Table 2. Newly-designed FISH probe (Seep1g-1443) and *nifH* mRNA HCR-FISH probe for labeling ANME-associated members of SEEP-SRB1g or SEEP-SRB1g *nifH* transcripts, respectively. Bolded sequence is complementary to HCR-FISH amplifier B1; nonbolded sequence is complementary to SEEP-SRB1g 16S rRNA or *nifH* RNA. Matches determined by comparison with ARB/SILVA SSU release 128 [54].

Supplemental Table 3. Stable isotope probing incubation conditions, sample sources and sulfide concentration measurements as a proxy for sulfate reduction activity.

Supplemental Table 4. SparCC-calculated correlations (pseudo-*p* < 0.01) between OTUs, detailing coefficients, OTU identifiers, and taxonomy assignments.

Supplemental Figure 1. Optimization of the newly designed Seep1g-1443 probe by FISH hybridization of ANME-2b—SEEP-SRB1g consortia at a range of formamide concentrations. Supplemental Figure 2. Krona chart depicting relative abundance of taxa in Costa Rica seep sediment sample #9279 (Fig. 4) as measured by 16S rRNA iTag amplicon sequencing.

Supplemental Figure 3. Krona chart depicting relative abundance of taxa in Costa Rica seep sediment sample #9112 (Fig. 4) as measured by 16S rRNA iTag amplicon sequencing.

Supplemental Figure 4. Krona chart depicting relative abundance of taxa in Costa Rica seep sediment sample #10073 (Fig. 7) as measured by 16S rRNA amplicon sequencing.

Supplemental Figure 5. 16S rRNA phylogeny inferred from maximum-likelihood methods using only full-length 16S rRNA sequences. Tree topology shown here is congruent with the phylogeny shown in Figure 3 constructed using a mix of shorter iTag amplicon and full-length 16S sequences.

Supplemental Figure 6. Extended *nifH* tree including unpublished SEEP-SRB1a MAGs possessing *nifH* sequences nearly identical to some recovered in environmental cDNA libraries (Dekas, et al. 2016).

Supplemental Figure 7. Negative controls for HCR-FISH experiments, demonstrating absence of signal in AOM aggregates with either only initiator or only amplifier added to HCR-FISH reactions.

Supplemental File 1. FASTA file containing the translated amino acid sequences for *nifH* included in Figure 7 in select ANME and SRB genomes (Chadwick, et al., *in prep*) and transcripts [8].

