## Supplemental Materials and Methods for "Experimentally-validated correlation analysis reveals new anaerobic methane oxidation partnerships with consortium-level heterogeneity in diazotrophy"

Supplementary Materials and Methods

*Sample collection*

Pushcore samples of seafloor sediment were collected by DSV *Alvin* during the May 20-June 11 2017 ROC HITS Expedition (AT37-13) aboard R/V *Atlantis* (operated by Woods Hole Oceanographic Institute, Woods Hole, MA, USA) to methane seep sites southwest of Costa Rica [1–3]. After retrieval from the seafloor, sediment pushcores were extruded aboard R/V *Atlantis* and sectioned at 1-3 cm intervals for geochemistry and microbiological sampling using published protocols [4, 5]. Subsamples for DNA extraction and microscopy were recovered using sterile cutoff 1 mL syringes (BD, Franklin Lakes, NJ, USA). Samples of seep carbonates and xenophyophores collected proximal to seafloor seep sites were also used for DNA extraction. Sediment, seep carbonate, and xenophyophore samples for DNA extraction were immediately frozen in liquid N_2_ and stored at -80˚C. Samples for microscopy were fixed in a filter-sterile (0.2 µm) 3X phosphate-buffered saline solution, pH 7.4 (145 mM NaCl, 1.4 mM NaH_2_PO_4_, 8 mM Na_2_HPO_4_, Sigma-Aldrich Corporation, St. Louis, MO, USA) with 2% paraformaldehyde (Electron Microscopy Sciences, Hatfield, PA, USA) for 24 h at 4˚C. Samples were subsequently washed with 1X PBS after centrifugation at 10000xg for 2 min. 1X PBS wash was removed after a second centrifugation and the resulting sediment pellet was resuspended in a 50:50 solution of ethanol and 1X PBS solution and stored at -20˚C.

Remaining sediment not used for DNA extraction or microscopy was placed in Mylar bags with filtered seawater, sparged with Ar, and stored at 4˚C after pressurization to ~2 atm with CH_4_. Upon return to the laboratory, Mylar bags were unsealed and decanted into 1L Pyrex bottles (Corning Life Sciences, Tewksbury, MA, USA) while being sparged with N_2_. Bottles were sealed with butyl stoppers and pressurized to ~2 atm with CH_4_. These incubations were stored in the dark at 4˚C for 1.5 yr before sampling, with spent media replaced every 3 months with fresh N_2_-sparged filter-sterilized seawater and methane. Mud from these incubations was also sampled for FISH and HCR-FISH by fixation in 4% paraformaldehyde at room temperature for 30 min. A full list of samples used in this study can be found in Supplementary Table 1.

*DNA Extraction and Illumina MiSeq sequencing of 16S rRNA gene*

DNA was extracted from 310 samples of Costa Rican methane seep sediments and seep carbonates (Supp. Table 1) using the Power Soil DNA Isolation Kit 12888 following manufacturer (Qiagen, Germantown, MD, USA) directions modified for sediment and carbonate samples [4, 6]. The V4-V5 region of the 16S rRNA gene was amplified using archaeal/bacterial primers [7] with Illumina (San Diego, CA, USA) adapters on 5’ end (515F: 5’-TCGTCGGCAGCGTCAGATGTGTATAAGAGACAG-GTGYCAGCMGCCGCGGTAA-3’, 926R: 5’-GTCTCGTGGGCTCGGAGATGTGTATAAGAGACAG-CCGYCAATTYMTTTRAGTTT-3’). PCR reaction mix was set up in duplicate for each sample with Q5 Hot Start High-Fidelity 2x Master Mix (New England Biolabs, Ipswich, MA, USA) in a 15 µL reaction volume according to manufacturer’s directions with annealing conditions of 54°C for 30 cycles. Duplicate PCR samples were then pooled and 2.5 µL of each product was barcoded with Illumina NexteraXT index 2 Primers that include unique 8-bp barcodes (P5 5’-AATGATACGGCGACCACCGAGATCTACAC-XXXXXXXX-TCGTCGGCAGCGTC-3’ and P7 5’-CAAGCAGAAGACGGCATACGAGAT-XXXXXXXX-GTCTCGTGGGCTCGG-3’). Amplification with barcoded primers used the same conditions as above, except for a volume of 25 µL, annealing at 66°C and 10 cycles. Products were purified using Millipore-Sigma (St. Louis, MO, USA) MultiScreen Plate MSNU03010 with vacuum manifold and quantified using ThermoFisherScientific (Waltham, MA, USA) QuantIT PicoGreen dsDNA Assay Kit P11496 on the BioRad CFX96 Touch Real-Time PCR Detection System. Barcoded samples were combined in equimolar amounts into single tube and purified with Qiagen PCR Purification Kit 28104 before submission to Laragen (Culver City, CA, USA) for 2 x 250 bp paired end analysis on Illumina’s MiSeq platform with PhiX addition of 15-20%.

*Processing of 16S rRNA gene MiSeq sequences*

Sequence data was processed in QIIME version 1.8.0 [8] following Mason, et al. 2015 [9]. Raw sequence pairs were joined and quality-trimmed using the default parameters in QIIME. Sequences were clustered into *de novo* operational taxonomic units (OTUs) with 99% similarity using UCLUST open reference clustering protocol and the most abundant sequence was chosen as representative for each *de novo* OTU [10]. Taxonomic identification for each representative sequence was assigned using the Silva-119 database [11] clustered at 99% similarity. This SILVA database had been appended with 1,197 in-house high-quality, methane seep-derived bacterial and archaeal full-length 16S rRNA sequences. Any sequences with pintail values > 75 were removed. The modified SILVA database is available upon request from the corresponding authors. Further taxonomic assignment of OTUs assigned to the SEEP-SRB1 clade was performed by aligning these 411 bp amplicon sequences to the Silva 119 database in ARB [12] and construction of a phylogenetic tree from full-length and amplicon 16S rRNA sequences of SEEP-SRB1 and sister clades to delineate SEEP-SRB1 subgroups [13]. Known contaminants in PCR reagents as determined by analysis of negative controls run with each MiSeq set were also removed [14] along with rare OTUs not present in any given library at a level of at least 10 reads.

*Correlation analysis of 16S rRNA amplicon libraries*

A QIIME-produced table of OTUs detected in the 310 methane seep sediment and seep carbonate amplicon libraries was further analyzed using the correlation algorithm SparCC [15]. A bash shell script (sparccWrapper.sh, written by Karoline Faust) was used to call SparCC Python scripts SparCC.py, MakeBootstraps.py, and PseudoPvals.py. First, SparCC.py calculated correlations between OTUs. MakeBootstraps.py then produced 100 shuffled OTU tables by random sampling from the real data with replacement and SparCC.py was used to calculate correlations in each of these 100 shuffled OTU tables. Finally, PseudoPvals.py calculated pseudo-*p* values for OTU correlations in the real dataset by comparison to correlations calculated in the shuffled OTU tables. As described by Friedman and Alm, 2012 [15], pseudo-*p-*values represent the fraction of correlation coefficients for a given pair of OTUs calculated from the 100 shuffled datasets that are greater than that calculated from the real datasets. Thus, a *pseudo*-p-values < 0.01 for a given pair of OTUs indicates that no correlation coefficient from any given shuffled dataset was greater than that calculated from our real data. Subsequent analysis of the produced tables describing magnitude and significance for OTU correlations was performed in R versions 3.3.3 and 3.5.0 [16], using visualization packages igraph [17], circlize [18], ggplot2 [19], and RColorBrewer [20]. In this study, only positive correlations (correlation coefficient > 0) between OTUs were used to examine potential ANME-SRB pairings. Analysis of cohesive blocks of OTUs (represented as nodes) in a force-directed network diagram [17, 21, 22] calculated from a filtered table of OTU correlations was interpreted to generate hypotheses of ANME-SRB pairings.

*Phylogenetic analysis of 16S rRNA amplicon sequences*

To examine phylogenetic placement of SRB 16S rRNA gene amplicon sequences predicted by network analysis to associate with particular ANME subgroup amplicon sequences, a phylogeny was constructed using RAxML-HPC [23] on XSEDE [24] using the CIPRES Science Gateway [25] from full-length 16S rRNA sequences of Deltaproteobacteria aligned by MUSCLE [26]. Although amplicon sequences contain significantly less information than full-length 16S rRNA sequences, they were used in phylogeny construction to allow direct comparison between amplicon and full length 16S sequences. 16S rRNA sequences were sourced from NCBI for published full-length 16S sequences [27], from 99% consensus OTU sequences produced by QIIME from amplicon libraries prepared from methane seep sediments (this work) and from genome contig files downloaded from the US Department of Energy Joint Genome Institute’s Integrated Microbial Genomes and Microbiomes (IMG/M) [72] of individual ANME-SRB consortia isolated by fluorescence-activated cell sorting of BONCAT-labeled consortia (BONCAT-FACS [27]). The latter was acquired either by direct download of 16S rRNA genes detected in genome bins or by tblastn (e-value < 1^-10^) searches of genome contig files using the 16S rRNA sequence from genome *Desulfosarcina* sp. BuS5 (IMG Genome ID 2513237157), closely related to known SEEP-SRB1a [28], as query. RAxML was run in parallel using raxmlHPC-HYBRID with the following settings: 100 bootstraps, 25 distinct rate categories, bootstrapping model GTRCAT, rapid bootstrapping, random seed for parsimony and for rapid bootstrapping set to 12345, and the Lewis ascertainment bias correction (called as raxmlHPC-HYBRID -T 4 -n result -s infile.txt -c 25 -m GTRCAT -p 12345 -k -f a -N 100 -x 12345 --asc-corr lewis). The resulting tree was exported and visualized using iTOL [29].

*FISH probe design for new ANME-associated SRB group SEEP-SRB1g*

A new FISH probe was designed in ARB using a modified version of the Silva 132 database (available on request). This new probe, named S-F-SP1g-1443-a-A-23 following published conventions [30] and hereafter referred to as Seep1g-1443 (5’-CCTCTCGCATAAAGCGAGTTAGC-3’, Supp. Table 2), was designed to complement and target 16S rRNA sequences in a monophyletic “*Desulfococcus* sp.” clade, which, based on phylogenetic analysis (see below), was renamed SEEP-SRB1g. Seep1g-1443 was ordered from Integrated DNA Technologies (Coralville, IA, USA) with fluor-dye Alexa488 attached to the 5’ end, prepared for use by dilution to 50 ng/µL, and frozen at -20˚C. FISH reaction conditions were optimized for Seep1g-1443 by performing a series of FISH reactions at a range of formamide concentrations between 20% to 45% vol/vol. In this range, signal was specific to the SRB partner with little observed cross-hybridization; optimal intensity and specificity at 35% (Supp. Fig. 1).

*FISH sediment sample preparation and imaging*

FISH and hybridization chain reaction (HCR-) FISH was performed on paraformaldehyde-fixed samples ANME-SRB consortia extracted from Costa Rican methane seep sediments using previously published density separation and FISH protocols [31]. Two samples of fixed sediment with abundant 16S iTAG amplicon reads of ANME-2a and -2b (sample 9279) or ANME-2c and -2b (sample 9112) were prepared for downstream FISH labeling and microscopy (Supp. Fig. 2). For each sample, 50 µL of fixed sediment was diluted with 950 µL 0.2 µm filter-sterilized 1X PBS in a 2 mL Eppendorf tube. After cooling for 10 min on ice, the diluted sediment was sonicated using a Branson Sonifier 150 (Branson Ultrasonics Corporation, Danbury, CT, USA). Sonication was performed with three 10 s pulses of the sonicator, set at 4 W output, with 10 s intervals between pulses. The 1 mL of sonicated sediment slurry was then pipetted onto 500 µL of Percoll (Sigma-Aldrich) and centrifuged at 16100 x G for 20 min at 4˚C. The supernatant with consortia was recovered and pipetted into 250 mL filter-sterile 1X PBS in a filter tower. This solution was filtered through a 5 µm polyethersulfone (PES) filter until ~50 mL solution remained in the tower. The filter was then washed with 200 mL 1X PBS while on the filter tower. Washing the remaining sample was repeated three times, with the final filtration step yielding a 1 mL aliquot. This 1mL aliquot was slowly concentrated onto a 0.2 µm GTTP white polycarbonate filter (Millipore-Sigma), keeping the filtered sample within a circular area of 0.5 mm-diameter. This area of the filter was then cut out with a razor blade and placed in a 250 µL PCR tube for FISH labeling.

FISH was performed overnight (18 hr) using the following modifications (G. Chadwick, pers. comm.) to previously-published protocols [27, 32]. A hybridization buffer at appropriate stringency was prepared along with accompanying wash buffer [33, 34] and pre-warmed to 46˚C and 48˚C in a hybridization oven and a water bath, respectively. 5 µL each of FISH probe stocks (50 ng/µL) Seep1g-1443 (this work), Seep1a-1441 [13], ANME-2a-828 (M. Aoki, pers. comm., 5’-GGTCGCACCGTGTCTGACACCT-3’), ANME-2b-729 [27], and ANME-2c-760 [35]. Four FISH experiments were performed, in which 5 µL each of 3 FISH probe stocks (at concentration 50 ng/µL) were added to 35 µL hybridization buffer in 200 µL PCR tubes along with the filter sections. Two experiments were performed at 20% formamide stringency on sample 9279, both using Seep1g-1443 (Alexa488) and Seep1a-1441 (cy5), and ether ANME-2b-729 (cy3) or ANME-2a-828 (cy3). Two similar experiments were performed at 45% formamide stringency on sample 9112 using instead either ANME-2b-729 (cy3) or ANME-2c-760 (cy3) and both Seep1g-1443 (Alexa488) and Seep1a-1441 (cy5). After 18 hr hybridization, filters were removed and incubated for 20 min at 48˚C in 200 µL wash buffer. Filter sections were then removed and briefly dipped in deionized water and placed on Superfrost Plus slides (Thermo Fischer Scientific, Waltham, MA, USA) to dry at room temperature in the dark. 10 µL 4,6-diamidino-2-phenylindole (DAPI, Sigma-Aldrich) dissolved in Citifluor (Electron Microscopy Sciences) was applied to filter sections and left to incubate for 15 min in the dark. A cover slip (No. 1.5, VWR, Radnor, PA, USA) was then placed on each filter.

Structured-illumination microscopy (SIM) was performed on FISH and HCR-FISH (see below) experiments to image ANME-SRB consortia at resolutions beyond that of traditional microscopy. After Immersol 518F immersion oil (Zeiss, Jena, Germany) was placed onto sample cover slips, FISH-labeled samples were examined using a Zeiss Elyra PS.1 SIM platform. Samples illuminated by Elyra laser lines (405 nm, 488 nm, 561 nm, 642 nm) and viewed through an alpha Plan-APOCHROMAT 100X/1.46 Oil DIC M27 objective and filter set (BP420-480+LP750, BP495-550+LP750, BP570-620+LP750, LP655) were imaged using a pco.edge sCMOS camera (PCO, Kelheim, Germany). Zen Black software (Zeiss) was used to construct final images from structured-illumination data.

*Imaging of nifH mRNA by HCR-FISH*

Hybridization chain reaction FISH (HCR-FISH) is a powerful technique to amplify signal from bound FISH probes by inducing polymerization of additional fluorophores to the bound probes [36, 37]. The protocol was modified from Yamaguchi and coworkers [38] and adapted to use lower probe concentrations (50 nM vs. 500 nM) and amplifier (300 nM) concentrations. In contrast to the published protocol, here, HCR-FISH was performed on white polycarbonate filters rather than directly on glass slides. HCR-FISH was performed using the same filter preparation protocol described above. This hybridization mix also included 5 µL each of 16S rRNA-targeted FISH probes Seep1g-1443 and ANME-2b-729 and a mix of HCR-FISH initiator probes (final concentration 50 nM) in the modified hybridization buffer (35% formamide stringency: 40 µL of 1M TRIS at pH 8 , 360 µL of 5M NaCL, 10 µL of 10% SDS, 700 µL of 100% formamide, 400 µL of 50% dextran sulfate, 4 µL of 50X Denhardt’s Solution, 486 µL of deionized water) designed to target SEEP-SRB1g *nifH* mRNA transcripts (Supp. Table 2). After 18 hr hybridization at 46˚C, filters were removed and placed in 200 µL wash buffer (4 µL 1M pH 8 TRIS, 3.2 µL 5M NaCl, 1 µL 10% SDS, 191.8 µL deionized water). Immediately after, an amplification buffer solution was prepared (200 µL 0.5 M NaH_2_PO_4_, 360 µL 5M NaCl, 2 µL 10% SDS, 400 µL 50% dextran sulfate, 4 µL 50X Denhardt’s Solution, 1034 µL deionized water). 5 µL each of hairpins B1H1 and B1H2 (3 µM stock) with attached Alexa647 fluorophores (Molecular Technologies, Pasadena, CA, USA) were added separately to two 45 µL volumes of amplification buffer in PCR tubes and snap cooled by placement in a C1000 Touch Thermal Cycler (BioRad, Hercules, CA, USA) for 3 min at 95˚C. Hairpins in amplification buffer were then left to cool at room temperature for 30 min. After the elapsed time, hairpins in amplification buffer were mixed and placed in PCR tubes. Filters were removed from wash buffer and placed in the mixed amplification buffer, and amplification was performed by placement of PCR tubes in a 35˚C water bath. After 15 min, filters were removed and placed into pre-chilled 1X PBS at 4˚C for 10 min. Filters were then removed and dipped in deionized water briefly before placement on Superfrost Plus slides to dry at room temperature in the dark. 10 µL DAPI in Citifluor was applied and No. 1.5 VWR coverslips were placed on filters. The HCR-FISH reaction with *nifH* probes was also performed in accordance with published protocols [39]. HCR-FISH v3.0 uses a different buffer system and longer incubation times during hybridization and amplification stages of the protocol but we observed similar results with both protocols.

*Comparative genomics of SEEP-SRB1g*

Genomes downloaded from the IMG/M database were searched using tblastn (e-value<1^-10^) for sequences matching reference NifD (NCBI Accession WP012698833), NifK (WP012698832), AprA (WP027353074), and DsrB (WP027352568) sequences. A reference sequence for chlorophyllide reductase BchX (WP011566468) was used as a reference sequence for a tblastn *nifH* search using BLAST+ on the command line [40]. The *nifH* search also included a set of cDNA sequences cloned from methane seep sediments using primers specific to *nifH* [40]. Phylogenetic trees of MUSCLE-aligned tblastn hits were calculated using RAxML on XSEDE through the CIPRES Science Gateway, using the following settings for RAxML: raxmlHPC-HYBRID_8.2.12_comet -n result -s infile.txt -c 25 -p 12345 -m PROTCATDAYHOFF -k -f a -N 100 -x 12345 --asc-corr lewis. Output was viewed in iTOL.

Genome trees were constructed using the Anvi’o platform [41] using HMM profiles from a subset of sequences from Campbell, et al. [42] consisting of only ribosomal proteins. HMM hits to these profiles were then concatenated, aligned in MUSCLE, and used as input in RAxML to generate genome trees (called with identical settings as those for individual gene trees).

*Stable isotope probing incubations with ^15^N_2_*

Incubated Costa Rica methane seep sediments from samples with abundant ANME-2b and SEEP-SRB1g (Supp. Fig. 4) were maintained in the laboratory under conditions supporting AOM and subsequently subsampled to test for diazotrophic activity in SEEP-SRB1g by stable isotope probing (SIP). SIP incubations (Supp. Table 3) were prepared by sparging source bottles and 30 mL serum bottles with N_2_ and mixing 5 mL of sediment with 5 mL N_2_-sparged artificial seawater without a N source (per L, 9.474 g MgCl_2_ • 6H_2_O, 0.2 g CaCl_2_ • 2H_2_O, 26.7 g NaCl, 0.522 g KCl, 1.42 g Na_2_SO_4_, 0.174 g K_2_HPO_4_, 1 mL L1 trace elements solution, 100 mL 250 mM pH 7.5 HEPES, 5 mL 1M NaHCO_3_, from a published medium composition [43]). Bottles were capped with butyl stoppers and overpressurized with CH_4_. Over the course of three days, 9 mL of artificial seawater supernatant was removed and replaced with 9 mL additional artificial seawater to remove residual NH_4_^+^_(aq)_. After pressurization to 2.8 bar CH_4_, two incubations were further pressurized with 1.2 mL ^15^N_2_ at 1 bar, approximately equivalent to 2% headspace in 20 mL CH_4_ at 2.8 bar. Two positive control incubations were inoculated with 20 µL 500 mM ^15^NH_4_Cl (^15^NH_4_Cl/NH_4_Cl = 0.1) and were further pressurized with 1.2 mL natural-abundance N_2_ at 1 bar. Incubations were sampled for microbial community analysis and geochemistry and refreshed every 3 months and samples for nanoSIMS were recovered after 9 months. Sulfate reduction activity was assayed using the published protocols [44].

*FISH-NanoSIMS*

Incubations were sampled for FISH-nanoSIMS [45] following fixation procedures described above. After fixation and Percoll separation, samples were embedded in 3% Difco Noble Agar (BD, USA) on a 5 µm polycarbonate filter, peeled off, dehydrated in an ethanol series, and embedded using Technovit H8100 Embedding kit (Kulzer GmbH, Wehrheim, Germany). 2 µm thin sections were cut using an Ultracut E microtome (Reichert AG, Wein, Austria) and mounted on Teflon/poly-L-lysine slides (Tekdon Inc., FL, USA) by placement on 50 µL H_2_O. FISH reactions were performed using Seep1g-1443 and ANME-2b-729 probes as described above, with the omission of 10% SDS to prevent detachment of section from slide (G. Chadwick, pers. comm.), and slides were imaged using a Zeiss Elyra PS.1 platform. After removal of DAPI-Citifuor by washing, slides were cut to fit into nanoSIMS sample holders and sputter-coated with 40 nm Au using a Cressington sputter coater. Spatially-resolved secondary-ion mass spectroscopy was then performed on sectioned ANME-SRB consortia using a Cameca NanoSIMS 50L housed in Caltech’s Microanalysis Center. Pre-sputtering of samples was performed using a 1 nA Cs^+^ ion beam until ^12^C^15^N^–^ ion counts stabilized. 512 x 512 pixel raster images of 20 µm^2^ were then collected for ^12^C^–^, ^16^O^–^, ^12^C^14^N^–^, ^15^N^12^C^–^, ^28^Si^–^, and ^32^S^–^ ions by sputtering with a ~1 pA primary Cs^+^ ion beam current with a dwell time of 12-48 ms/pixel. Mass calibration was performed once an hour for all masses. NanoSIMS data were processed using look@nanoSIMS [46] to determine ^15^N fractional abundance, ^15^N/(^15^N+^14^N). Regions of interest (ROIs) for ANME-2b and SEEP-SRB1g in consortia were drawn with Adobe Draw using secondary electron images of sectioned consortia compared with FISH images of the same section collected prior to nanoSIMS. ROIs annotated as ANME-2b or SEEP-SRB1g were then used as input for a MATLAB script used to extract ^15^N fractional abundance from ROIs.
