## Supplemental Figure 2 for "Experimentally-validated correlation analysis reveals new anaerobic methane oxidation partnerships with consortium-level heterogeneity in diazotrophy": Supplemental_Figure_2.html

Javascript must be enabled to view this page.

magnitude

 23514

 14247

 0

 0

 0

 0

 0

 0

 0

 0

 0

 0

 0

 0

 0

 0

 0

 0

 0

 0

 14214

 0

 0

 0

 0

 0

 0

 0

 0

 11

 11

 3

 0

 0

 3

 8

 8

 0

 0

 0

 0

 0

 0

 0

 0

 0

 0

 0

 0

 0

 0

 0

 0

 14198

 0

 0

 0

 0

 0

 0

 0

 0

 0

 0

 0

 0

 0

 0

 0

 0

 0

 0

 1

 0

 0

 1

 1

 0

 0

 0

 0

 0

 0

 0

 14197

 38

 38

 13913

 396

 10758

 2759

 105

 105

 0

 0

 0

 0

 0

 0

 141

 133

 8

 0

 0

 0

 0

 0

 0

 0

 0

 0

 0

 5

 0

 0

 0

 0

 0

 0

 0

 0

 0

 0

 0

 0

 0

 0

 0

 0

 0

 0

 5

 0

 0

 0

 0

 3

 3

 0

 0

 0

 0

 1

 1

 0

 0

 0

 0

 0

 0

 0

 0

 1

 1

 0

 0

 0

 0

 0

 0

 0

 0

 0

 0

 0

 0

 0

 0

 0

 0

 0

 0

 0

 33

 0

 0

 0

 0

 0

 0

 0

 0

 0

 0

 0

 0

 0

 0

 0

 0

 13

 13

 13

 13

 0

 0

 0

 0

 10

 10

 10

 10

 10

 3

 3

 3

 5

 5

 0

 0

 5

 2

 2

 2

 0

 0

 0

 0

 0

 0

 0

 0

 0

 0

 0

 0

 0

 0

 0

 0

 0

 0

 0

 0

 0

 0

 0

 0

 0

 0

 0

 0

 0

 0

 0

 0

 0

 0

 8117

 39

 39

 39

 39

 39

 894

 0

 0

 0

 0

 14

 0

 0

 0

 0

 0

 0

 0

 0

 0

 0

 0

 0

 0

 0

 0

 0

 0

 0

 0

 0

 0

 3

 3

 3

 0

 0

 0

 0

 0

 0

 6

 6

 6

 0

 0

 0

 0

 0

 0

 0

 0

 0

 0

 0

 0

 0

 0

 0

 0

 0

 0

 0

 0

 0

 0

 0

 0

 0

 0

 0

 0

 0

 0

 5

 5

 5

 0

 0

 0

 864

 0

 0

 0

 6

 6

 6

 0

 0

 0

 0

 0

 0

 0

 0

 0

 0

 13

 1

 1

 0

 0

 0

 0

 0

 0

 12

 12

 0

 0

 845

 845

 845

 0

 0

 0

 0

 0

 0

 16

 16

 16

 16

 0

 0

 0

 0

 56

 0

 0

 0

 0

 45

 45

 0

 0

 5

 3

 0

 2

 0

 0

 39

 39

 1

 1

 0

 0

 0

 0

 3

 1

 1

 1

 0

 0

 0

 0

 0

 0

 0

 0

 0

 0

 0

 0

 0

 0

 0

 0

 0

 0

 0

 0

 0

 0

 0

 0

 0

 0

 0

 0

 2

 0

 0

 0

 0

 0

 0

 0

 0

 0

 0

 2

 0

 2

 0

 0

 0

 0

 0

 0

 0

 0

 0

 0

 0

 0

 0

 0

 0

 0

 0

 0

 0

 0

 0

 0

 0

 0

 0

 0

 0

 0

 0

 0

 0

 0

 0

 0

 0

 0

 0

 0

 0

 0

 0

 0

 0

 0

 3

 3

 3

 3

 0

 0

 0

 0

 0

 0

 0

 0

 0

 0

 0

 0

 2

 2

 2

 2

 3

 0

 0

 0

 1

 0

 0

 0

 0

 1

 1

 2

 2

 2

 0

 0

 0

 0

 0

 0

 0

 0

 0

 0

 0

 0

 0

 0

 0

 0

 0

 0

 0

 0

 0

 0

 0

 0

 0

 0

 0

 0

 5

 0

 0

 0

 0

 5

 5

 5

 5

 0

 0

 0

 0

 0

 0

 0

 0

 0

 0

 1051

 118

 118

 118

 118

 115

 115

 115

 115

 0

 0

 0

 0

 0

 0

 0

 0

 19

 19

 0

 0

 0

 0

 0

 0

 19

 0

 0

 19

 0

 0

 0

 0

 0

 0

 0

 0

 0

 0

 0

 54

 54

 0

 0

 0

 0

 0

 0

 0

 0

 0

 0

 0

 0

 0

 54

 2

 0

 0

 0

 0

 0

 0

 0

 0

 23

 0

 29

 0

 0

 0

 0

 0

 0

 0

 292

 292

 1

 1

 0

 0

 30

 4

 0

 0

 0

 1

 0

 25

 261

 213

 0

 0

 0

 0

 0

 0

 0

 0

 0

 0

 11

 0

 0

 10

 0

 0

 0

 0

 0

 0

 0

 0

 0

 0

 4

 23

 0

 0

 0

 0

 0

 0

 0

 0

 0

 0

 6

 6

 6

 6

 54

 54

 54

 54

 72

 72

 10

 10

 0

 0

 0

 0

 0

 0

 0

 0

 0

 0

 0

 0

 0

 0

 0

 0

 0

 0

 0

 0

 0

 0

 8

 2

 0

 0

 4

 0

 0

 2

 0

 0

 0

 54

 54

 0

 0

 321

 321

 321

 321

 0

 0

 0

 0

 0

 0

 0

 0

 0

 0

 0

 0

 0

 0

 0

 0

 0

 0

 0

 0

 9

 9

 9

 9

 9

 9

 9

 9

 9

 9

 0

 0

 0

 0

 0

 4

 0

 0

 0

 0

 0

 0

 0

 0

 4

 4

 4

 4

 0

 0

 0

 0

 0

 0

 0

 0

 0

 0

 38

 38

 38

 38

 38

 0

 0

 0

 0

 0

 0

 0

 0

 0

 0

 372

 0

 0

 0

 0

 0

 0

 0

 0

 372

 372

 372

 372

 0

 0

 0

 0

 0

 0

 0

 0

 0

 0

 0

 0

 0

 0

 0

 0

 0

 0

 89

 3

 3

 3

 3

 0

 0

 86

 86

 11

 11

 0

 0

 0

 0

 0

 0

 64

 64

 11

 11

 307

 2

 2

 2

 2

 269

 269

 269

 1

 0

 0

 0

 0

 268

 17

 0

 0

 0

 17

 17

 17

 1

 1

 1

 0

 1

 0

 0

 0

 0

 0

 0

 5

 3

 3

 3

 0

 0

 0

 0

 0

 0

 0

 0

 0

 0

 0

 0

 0

 0

 0

 0

 0

 0

 0

 0

 0

 0

 0

 0

 0

 0

 0

 0

 2

 2

 2

 0

 0

 0

 0

 0

 0

 0

 0

 0

 0

 0

 0

 3

 3

 3

 3

 0

 0

 0

 0

 0

 0

 0

 0

 0

 0

 0

 0

 0

 0

 0

 0

 0

 0

 0

 0

 0

 2

 2

 2

 2

 0

 0

 0

 0

 0

 0

 0

 0

 0

 0

 0

 0

 0

 0

 0

 0

 0

 0

 0

 0

 0

 0

 0

 8

 8

 8

 8

 6

 0

 0

 0

 0

 2

 2

 2

 2

 4

 0

 0

 0

 4

 4

 0

 0

 0

 0

 4

 0

 0

 0

 0

 0

 0

 0

 0

 0

 0

 0

 0

 0

 0

 0

 0

 0

 0

 0

 0

 0

 0

 0

 0

 0

 0

 0

 14

 14

 14

 0

 0

 0

 0

 9

 9

 1

 1

 0

 0

 4

 4

 0

 0

 0

 0

 0

 0

 0

 0

 0

 0

 0

 0

 0

 0

 0

 0

 0

 0

 1

 1

 0

 0

 0

 0

 0

 0

 0

 0

 0

 1

 1

 1

 0

 0

 0

 0

 0

 0

 0

 0

 0

 0

 0

 0

 0

 0

 0

 0

 0

 0

 0

 0

 0

 2

 2

 0

 0

 0

 0

 0

 0

 0

 0

 0

 0

 0

 0

 0

 0

 0

 0

 0

 0

 0

 0

 0

 0

 0

 2

 2

 2

 119

 0

 0

 0

 0

 0

 0

 0

 0

 0

 0

 0

 0

 0

 0

 0

 0

 0

 0

 0

 0

 0

 0

 0

 0

 0

 0

 0

 0

 0

 0

 0

 0

 0

 0

 0

 0

 0

 0

 0

 0

 0

 0

 0

 0

 0

 0

 0

 0

 0

 0

 0

 0

 0

 0

 0

 0

 0

 0

 0

 0

 0

 0

 0

 0

 0

 119

 0

 0

 0

 119

 5

 5

 0

 0

 31

 2

 1

 28

 31

 18

 0

 0

 0

 13

 0

 0

 0

 0

 0

 0

 0

 0

 0

 0

 0

 0

 0

 0

 0

 0

 0

 0

 0

 0

 0

 0

 10

 0

 10

 0

 0

 0

 0

 0

 0

 0

 0

 0

 0

 0

 0

 0

 0

 0

 0

 2

 0

 0

 0

 0

 2

 0

 0

 1

 1

 1

 0

 0

 0

 0

 0

 0

 1

 16

 3

 9

 0

 4

 22

 4

 0

 0

 0

 18

 0

 0

 0

 0

 0

 0

 0

 0

 0

 0

 0

 0

 0

 0

 0

 0

 0

 0

 0

 0

 0

 0

 0

 0

 0

 0

 0

 0

 0

 0

 0

 0

 0

 0

 0

 0

 0

 0

 0

 0

 0

 0

 0

 0

 0

 0

 0

 0

 0

 2

 2

 2

 2

 0

 0

 0

 2

 0

 0

 0

 0

 0

 0

 0

 57

 57

 1

 1

 1

 10

 10

 10

 2

 2

 2

 0

 0

 0

 44

 44

 44

 0

 0

 0

 22

 22

 22

 22

 22

 2

 2

 2

 2

 2

 0

 0

 0

 0

 0

 36

 2

 2

 2

 2

 0

 0

 0

 0

 0

 0

 0

 0

 0

 0

 0

 0

 0

 0

 0

 0

 0

 0

 0

 0

 0

 0

 0

 0

 0

 0

 0

 0

 0

 0

 0

 0

 0

 0

 0

 2

 2

 2

 2

 0

 0

 0

 0

 0

 0

 0

 0

 0

 0

 0

 0

 0

 0

 28

 28

 28

 28

 0

 0

 0

 0

 0

 0

 0

 0

 4

 4

 4

 4

 0

 0

 0

 0

 0

 0

 0

 0

 0

 0

 0

 0

 0

 6

 6

 6

 0

 0

 0

 0

 6

 6

 0

 0

 0

 0

 0

 0

 0

 0

 0

 0

 0

 0

 0

 0

 0

 0

 0

 0

 0

 152

 0

 0

 0

 0

 0

 0

 0

 0

 0

 0

 0

 0

 0

 0

 0

 0

 0

 0

 0

 0

 12

 12

 12

 12

 64

 3

 3

 3

 0

 0

 0

 0

 0

 0

 0

 0

 0

 0

 0

 0

 0

 0

 0

 0

 0

 0

 0

 0

 0

 11

 11

 11

 32

 32

 32

 0

 0

 0

 13

 6

 6

 1

 1

 0

 0

 1

 1

 5

 0

 0

 1

 0

 0

 0

 1

 1

 2

 0

 0

 0

 0

 0

 0

 0

 4

 4

 4

 1

 1

 1

 0

 0

 0

 0

 0

 0

 0

 0

 0

 0

 0

 0

 0

 2

 2

 2

 2

 74

 0

 0

 0

 0

 0

 74

 74

 19

 5

 0

 0

 0

 0

 20

 0

 6

 5

 0

 0

 19

 0

 0

 0

 0

 0

 0

 0

 0

 0

 0

 0

 0

 4737

 47

 47

 47

 47

 0

 0

 0

 0

 0

 0

 0

 0

 0

 0

 0

 0

 75

 5

 5

 5

 0

 0

 0

 2

 0

 0

 2

 2

 0

 0

 0

 0

 0

 0

 0

 0

 0

 0

 0

 0

 0

 0

 0

 0

 0

 0

 0

 0

 0

 0

 0

 0

 31

 0

 0

 0

 0

 0

 0

 0

 0

 0

 0

 0

 0

 0

 0

 0

 0

 0

 0

 13

 0

 0

 1

 0

 0

 0

 0

 12

 0

 0

 0

 0

 0

 0

 0

 0

 0

 0

 4

 3

 0

 0

 0

 0

 0

 1

 0

 0

 0

 0

 0

 0

 0

 0

 0

 14

 0

 0

 0

 0

 14

 0

 0

 0

 0

 0

 0

 0

 0

 30

 30

 2

 0

 0

 0

 0

 5

 0

 0

 0

 0

 0

 0

 0

 0

 0

 0

 0

 0

 0

 0

 0

 0

 0

 0

 0

 0

 0

 0

 23

 7

 0

 0

 0

 0

 0

 0

 0

 0

 0

 0

 0

 0

 0

 0

 0

 0

 0

 0

 7

 0

 0

 0

 0

 0

 0

 0

 0

 0

 0

 0

 0

 0

 7

 0

 0

 0

 0

 0

 0

 0

 0

 0

 0

 0

 0

 0

 0

 0

 0

 0

 0

 0

 0

 0

 0

 0

 0

 0

 0

 0

 0

 0

 0

 0

 0

 0

 0

 0

 0

 0

 0

 0

 0

 0

 0

 0

 0

 0

 0

 0

 0

 0

 0

 0

 0

 0

 0

 0

 0

 0

 0

 0

 0

 0

 0

 0

 0

 0

 0

 0

 0

 0

 0

 0

 1

 0

 0

 0

 0

 0

 0

 0

 0

 0

 0

 0

 0

 0

 0

 0

 0

 0

 0

 0

 0

 0

 0

 0

 0

 0

 0

 0

 0

 0

 0

 0

 0

 0

 0

 1

 1

 0

 0

 1

 0

 0

 0

 0

 0

 0

 0

 0

 0

 0

 0

 0

 0

 0

 0

 0

 0

 0

 0

 0

 0

 0

 0

 0

 0

 2475

 32

 32

 32

 0

 0

 0

 5

 5

 2

 0

 0

 0

 2

 0

 1

 0

 0

 0

 0

 0

 0

 0

 0

 0

 146

 146

 0

 0

 146

 2184

 12

 12

 1183

 78

 0

 0

 0

 0

 0

 0

 85

 839

 0

 0

 0

 0

 4

 0

 0

 0

 0

 0

 101

 30

 0

 46

 986

 160

 138

 331

 0

 0

 2

 0

 0

 12

 0

 337

 0

 6

 3

 0

 0

 0

 3

 0

 0

 0

 0

 0

 0

 0

 0

 0

 0

 0

 0

 0

 0

 0

 0

 0

 0

 19

 1

 1

 0

 0

 0

 0

 0

 0

 3

 0

 0

 3

 0

 2

 2

 0

 0

 0

 0

 0

 0

 13

 13

 0

 0

 0

 0

 0

 0

 0

 2

 2

 2

 42

 3

 3

 0

 0

 0

 0

 0

 0

 0

 0

 0

 0

 0

 0

 0

 0

 0

 0

 0

 0

 0

 0

 0

 12

 0

 0

 12

 2

 2

 0

 0

 0

 0

 25

 25

 8

 8

 8

 22

 22

 22

 13

 13

 13

 2

 0

 0

 1

 0

 0

 1

 0

 0

 0

 1

 0

 0

 1

 0

 0

 0

 0

 528

 528

 0

 0

 0

 0

 0

 0

 0

 528

 5

 0

 184

 339

 0

 0

 0

 0

 0

 0

 0

 0

 1577

 263

 263

 263

 0

 0

 0

 0

 0

 0

 0

 0

 0

 0

 0

 17

 0

 0

 17

 10

 0

 0

 0

 0

 0

 0

 0

 0

 0

 0

 0

 0

 0

 7

 0

 0

 0

 0

 0

 0

 0

 0

 0

 0

 0

 0

 0

 0

 0

 0

 0

 0

 0

 0

 0

 0

 0

 0

 0

 0

 7

 7

 7

 0

 0

 0

 0

 0

 0

 108

 108

 108

 143

 2

 2

 3

 0

 3

 0

 0

 138

 0

 138

 0

 0

 0

 0

 0

 0

 0

 0

 0

 13

 13

 13

 0

 0

 0

 0

 0

 0

 0

 0

 0

 0

 0

 0

 0

 0

 0

 0

 0

 12

 12

 0

 0

 0

 0

 12

 0

 0

 0

 0

 0

 0

 0

 0

 2

 2

 0

 2

 0

 0

 0

 0

 0

 0

 0

 0

 40

 22

 22

 0

 0

 0

 0

 17

 17

 1

 1

 0

 0

 0

 0

 0

 0

 0

 0

 0

 0

 0

 0

 0

 0

 0

 0

 0

 0

 0

 0

 0

 0

 0

 0

 0

 0

 0

 0

 0

 0

 0

 0

 0

 0

 0

 0

 0

 0

 0

 0

 0

 0

 0

 0

 0

 0

 0

 0

 0

 0

 0

 0

 0

 0

 0

 0

 0

 0

 0

 0

 0

 0

 0

 0

 0

 0

 0

 0

 0

 0

 0

 0

 0

 0

 0

 0

 0

 0

 0

 0

 0

 649

 649

 649

 0

 0

 0

 0

 0

 0

 0

 0

 0

 0

 0

 0

 0

 0

 0

 0

 0

 0

 0

 8

 8

 8

 0

 0

 0

 269

 0

 0

 0

 0

 0

 0

 233

 21

 0

 10

 0

 0

 0

 198

 4

 36

 0

 0

 0

 0

 0

 2

 0

 0

 0

 34

 0

 0

 11

 11

 0

 0

 0

 0

 1

 10

 0

 0

 0

 0

 35

 4

 4

 31

 31

 0

 0

 0

 0

 0

 0

 0

 0

 0

 0

 4

 4

 4

 4

 0

 0

 0

 0

 30

 30

 30

 30

 0

 0

 0

 0

 0

 0

 0

 0

 0

 0

 0

 0

 0

 0

 0

 0

 0

 0

 0

 0

 0

 0

 0

 0

 0

 0

 0

 0

 0

 0

 0

 0

 0

 0

 0

 0

 0

 0

 0

 0

 0

 0

 0

 0

 0

 0

 0

 42

 42

 9

 9

 9

 2

 2

 2

 0

 0

 0

 0

 0

 0

 4

 4

 4

 0

 0

 0

 27

 0

 0

 0

 0

 0

 0

 0

 0

 0

 0

 0

 27

 0

 27

 0

 0

 0

 0

 0

 9

 9

 9

 9

 9

 10

 10

 10

 10

 10

 4

 4

 0

 0

 0

 0

 0

 0

 0

 0

 0

 0

 0

 0

 4

 4

 4

 0

 0

 0

 0

 0

 0

 0

 0

 0

 0

 23

 0

 0

 0

 0

 0

 0

 0

 0

 0

 0

 0

 0

 0

 0

 0

 3

 0

 0

 0

 3

 3

 3

 0

 0

 0

 0

 0

 0

 0

 0

 0

 0

 0

 0

 0

 0

 0

 0

 0

 0

 0

 0

 0

 0

 0

 0

 0

 0

 0

 0

 0

 0

 0

 0

 0

 0

 0

 20

 20

 4

 4

 1

 1

 0

 0

 1

 1

 14

 0

 0

 14

 0

 0

 0

 0

 0

 0

 0

 0

 0

 0

 0

 0

 0

 0

 0

 0

 0

 0

 0

 0

 0

 0

 0

 0

 0

 0

 0

 0

 0

 0

 0

 0

 0

 0

 0

 0

 0

 0

 0

 0

 0

 0

 0

 0

 0

 0

 0

 0

 0

 0

 1150

 1150

 1150

 1150

 1150

 1150
