## Supplemental Figure 3 for "Experimentally-validated correlation analysis reveals new anaerobic methane oxidation partnerships with consortium-level heterogeneity in diazotrophy": Supplemental_Figure_3.html

Javascript must be enabled to view this page.

magnitude

 27984

 15034

 0

 0

 0

 0

 0

 6

 6

 6

 6

 6

 0

 0

 0

 0

 0

 0

 0

 0

 14868

 0

 0

 0

 0

 0

 0

 0

 0

 58

 58

 4

 0

 0

 4

 34

 34

 0

 0

 18

 18

 2

 0

 2

 0

 0

 0

 0

 0

 0

 0

 0

 0

 14658

 0

 0

 0

 3825

 60

 60

 2227

 2227

 1487

 1487

 51

 51

 0

 0

 0

 0

 0

 0

 1

 0

 0

 1

 1

 0

 0

 0

 0

 0

 0

 0

 10832

 367

 367

 3321

 99

 2356

 866

 7033

 7033

 2

 2

 0

 0

 15

 15

 94

 28

 66

 0

 0

 0

 0

 0

 0

 0

 0

 0

 0

 152

 0

 0

 0

 2

 2

 2

 0

 0

 0

 2

 2

 2

 0

 0

 0

 0

 0

 0

 148

 0

 0

 0

 0

 9

 9

 0

 0

 0

 0

 38

 38

 0

 0

 0

 0

 91

 91

 0

 0

 0

 0

 0

 0

 0

 0

 10

 10

 0

 0

 0

 0

 0

 0

 0

 0

 0

 0

 0

 0

 0

 160

 0

 0

 0

 0

 0

 0

 0

 0

 0

 0

 0

 0

 0

 0

 0

 0

 41

 41

 41

 41

 0

 0

 0

 0

 114

 114

 114

 114

 4

 0

 0

 0

 2

 2

 0

 0

 2

 2

 2

 2

 1

 1

 1

 1

 0

 0

 0

 0

 0

 0

 0

 0

 0

 0

 0

 0

 0

 0

 0

 0

 0

 0

 0

 0

 0

 0

 0

 0

 0

 0

 0

 0

 0

 0

 11073

 67

 67

 67

 67

 67

 106

 0

 0

 0

 0

 5

 0

 0

 0

 0

 0

 0

 0

 0

 0

 0

 0

 0

 0

 0

 0

 0

 0

 0

 0

 0

 0

 0

 0

 0

 0

 0

 0

 0

 0

 0

 4

 4

 4

 0

 0

 0

 0

 0

 0

 0

 0

 0

 0

 0

 0

 0

 0

 0

 0

 0

 0

 0

 0

 0

 0

 0

 0

 0

 0

 0

 0

 0

 0

 1

 1

 1

 0

 0

 0

 81

 0

 0

 0

 7

 7

 7

 0

 0

 0

 0

 0

 0

 0

 0

 0

 0

 1

 0

 0

 0

 0

 0

 0

 0

 0

 1

 1

 0

 0

 72

 72

 72

 0

 0

 0

 1

 1

 1

 18

 18

 18

 18

 2

 2

 2

 2

 46

 0

 0

 0

 0

 30

 30

 0

 0

 1

 0

 0

 1

 0

 0

 29

 29

 0

 0

 0

 0

 0

 0

 0

 0

 0

 0

 0

 0

 0

 0

 0

 0

 0

 0

 0

 0

 0

 0

 0

 0

 0

 0

 0

 0

 0

 0

 0

 0

 0

 0

 0

 0

 0

 0

 0

 0

 0

 0

 0

 0

 0

 0

 0

 0

 0

 0

 0

 0

 0

 0

 0

 0

 0

 0

 0

 0

 0

 0

 0

 0

 0

 0

 0

 0

 0

 0

 0

 0

 0

 0

 0

 0

 0

 0

 0

 0

 0

 0

 0

 0

 0

 0

 0

 0

 0

 0

 0

 0

 0

 0

 0

 0

 4

 4

 4

 4

 0

 0

 0

 0

 0

 0

 0

 0

 0

 0

 0

 0

 12

 12

 12

 12

 0

 0

 0

 0

 0

 0

 0

 0

 0

 0

 0

 0

 0

 0

 0

 0

 0

 0

 0

 0

 0

 0

 0

 0

 0

 0

 0

 0

 0

 0

 0

 0

 0

 0

 0

 0

 0

 0

 0

 0

 0

 0

 0

 0

 0

 0

 0

 0

 0

 0

 0

 4

 4

 4

 4

 4

 1

 1

 1

 1

 1

 1374

 152

 152

 152

 152

 148

 148

 148

 148

 0

 0

 0

 0

 0

 0

 0

 0

 37

 37

 0

 0

 0

 0

 0

 0

 37

 0

 0

 37

 0

 0

 0

 0

 0

 0

 0

 0

 0

 0

 0

 20

 20

 0

 0

 0

 0

 0

 0

 0

 0

 0

 0

 0

 0

 0

 20

 1

 0

 0

 0

 0

 0

 0

 0

 0

 4

 0

 15

 0

 0

 0

 0

 0

 0

 0

 121

 121

 2

 2

 0

 0

 12

 1

 0

 0

 0

 0

 0

 11

 107

 56

 0

 0

 0

 0

 0

 0

 0

 0

 0

 0

 3

 0

 0

 20

 0

 0

 0

 0

 0

 0

 0

 0

 0

 0

 0

 28

 0

 0

 0

 0

 0

 0

 0

 0

 0

 0

 83

 83

 83

 83

 62

 62

 62

 62

 224

 224

 47

 47

 0

 0

 1

 0

 0

 0

 0

 1

 0

 0

 0

 0

 0

 0

 0

 0

 0

 0

 0

 0

 0

 0

 5

 0

 0

 0

 0

 0

 0

 5

 0

 0

 0

 171

 171

 0

 0

 527

 527

 527

 527

 0

 0

 0

 0

 0

 0

 0

 0

 0

 0

 0

 0

 0

 0

 0

 0

 0

 0

 0

 0

 10

 10

 10

 10

 10

 579

 579

 579

 579

 579

 0

 0

 0

 0

 0

 0

 0

 0

 0

 0

 0

 0

 0

 0

 0

 0

 0

 0

 0

 0

 0

 0

 0

 13

 13

 13

 13

 13

 43

 43

 43

 43

 43

 0

 0

 0

 0

 0

 3

 3

 3

 3

 3

 429

 2

 2

 2

 2

 0

 0

 0

 0

 427

 427

 427

 427

 1

 1

 1

 0

 0

 0

 0

 0

 0

 0

 0

 0

 0

 0

 1

 1

 0

 0

 73

 13

 13

 12

 12

 1

 1

 60

 60

 2

 2

 8

 8

 0

 0

 0

 0

 24

 24

 26

 26

 580

 11

 11

 11

 11

 508

 508

 508

 1

 0

 0

 0

 0

 507

 13

 0

 0

 0

 13

 13

 13

 0

 0

 0

 0

 0

 0

 0

 0

 0

 0

 0

 25

 6

 6

 6

 0

 0

 0

 0

 0

 0

 0

 0

 0

 0

 0

 0

 0

 7

 7

 7

 7

 7

 7

 2

 2

 2

 0

 0

 0

 3

 3

 3

 0

 0

 0

 0

 0

 0

 0

 0

 0

 0

 0

 0

 0

 0

 0

 2

 2

 2

 2

 0

 0

 0

 0

 0

 0

 0

 0

 0

 0

 0

 0

 0

 0

 0

 0

 0

 0

 0

 0

 0

 0

 0

 0

 0

 0

 0

 0

 0

 0

 0

 0

 0

 0

 0

 0

 0

 0

 0

 0

 0

 0

 0

 0

 0

 0

 0

 0

 21

 21

 21

 21

 1

 0

 0

 0

 0

 0

 0

 0

 0

 1

 0

 0

 0

 1

 1

 0

 0

 0

 0

 1

 0

 0

 0

 0

 0

 0

 0

 0

 0

 0

 0

 0

 0

 0

 0

 0

 0

 0

 0

 0

 0

 0

 0

 0

 0

 0

 0

 238

 238

 238

 0

 0

 0

 0

 217

 217

 2

 2

 3

 3

 16

 16

 0

 0

 0

 0

 0

 0

 0

 0

 0

 0

 0

 0

 0

 0

 0

 0

 0

 0

 9

 9

 0

 0

 0

 0

 0

 0

 0

 0

 0

 4

 4

 4

 0

 0

 0

 3

 3

 3

 2

 2

 2

 0

 0

 0

 0

 0

 0

 0

 0

 0

 0

 0

 0

 27

 27

 1

 1

 1

 0

 0

 0

 0

 0

 0

 0

 0

 0

 0

 0

 0

 0

 0

 0

 0

 0

 0

 0

 0

 26

 26

 26

 23

 0

 0

 0

 0

 0

 0

 0

 0

 0

 0

 0

 0

 0

 0

 0

 0

 0

 0

 0

 0

 0

 0

 0

 0

 0

 0

 0

 0

 0

 0

 0

 0

 0

 0

 0

 0

 0

 0

 0

 0

 0

 0

 0

 0

 0

 0

 0

 0

 0

 0

 0

 0

 0

 0

 0

 0

 0

 0

 0

 0

 0

 0

 0

 0

 0

 23

 0

 0

 0

 23

 5

 5

 0

 0

 3

 0

 0

 3

 0

 0

 0

 0

 0

 0

 0

 0

 0

 0

 0

 0

 0

 0

 0

 0

 0

 0

 0

 0

 0

 0

 0

 0

 0

 0

 0

 0

 2

 0

 2

 0

 0

 0

 0

 0

 0

 0

 0

 0

 0

 0

 0

 0

 0

 0

 0

 0

 0

 0

 0

 0

 0

 0

 0

 0

 0

 0

 0

 0

 0

 0

 0

 0

 0

 0

 0

 0

 0

 0

 13

 1

 0

 0

 0

 12

 0

 0

 0

 0

 0

 0

 0

 0

 0

 0

 0

 0

 0

 0

 0

 0

 0

 0

 0

 0

 0

 0

 0

 0

 0

 0

 0

 0

 0

 0

 0

 0

 0

 0

 0

 0

 0

 0

 0

 0

 0

 0

 0

 0

 0

 0

 0

 0

 0

 2

 2

 2

 2

 0

 0

 0

 2

 0

 0

 8

 8

 8

 8

 8

 7

 7

 0

 0

 0

 4

 4

 4

 1

 1

 1

 0

 0

 0

 2

 2

 2

 0

 0

 0

 149

 149

 149

 149

 149

 4

 4

 4

 4

 4

 0

 0

 0

 0

 0

 37

 6

 6

 6

 6

 0

 0

 0

 0

 0

 0

 0

 0

 0

 0

 0

 0

 1

 1

 1

 1

 0

 0

 0

 0

 0

 0

 0

 0

 0

 0

 0

 0

 0

 0

 0

 0

 0

 0

 0

 1

 1

 1

 1

 5

 0

 0

 0

 1

 1

 1

 4

 4

 4

 0

 0

 0

 0

 21

 21

 21

 21

 0

 0

 0

 0

 0

 0

 0

 0

 3

 3

 3

 3

 0

 0

 0

 0

 0

 0

 0

 0

 1

 1

 1

 1

 1

 5

 5

 5

 0

 0

 0

 0

 4

 4

 0

 0

 0

 0

 1

 0

 0

 0

 0

 1

 0

 0

 0

 0

 0

 0

 0

 0

 0

 433

 0

 0

 0

 0

 0

 0

 0

 0

 2

 2

 2

 2

 0

 0

 0

 0

 0

 0

 0

 0

 25

 25

 25

 25

 297

 35

 35

 35

 0

 0

 0

 0

 0

 0

 0

 0

 0

 0

 0

 0

 4

 4

 4

 0

 0

 0

 1

 1

 1

 74

 74

 74

 57

 57

 57

 0

 0

 0

 114

 22

 22

 11

 11

 65

 65

 3

 3

 13

 0

 0

 0

 0

 4

 0

 1

 4

 3

 1

 0

 0

 0

 0

 0

 0

 0

 0

 0

 7

 7

 7

 2

 2

 2

 0

 0

 0

 3

 3

 3

 6

 6

 6

 6

 41

 41

 41

 41

 53

 0

 0

 0

 0

 0

 53

 53

 23

 0

 0

 0

 0

 0

 24

 0

 0

 1

 0

 0

 5

 0

 0

 0

 0

 0

 0

 0

 0

 9

 9

 9

 9

 6295

 38

 38

 38

 38

 0

 0

 0

 0

 0

 0

 0

 0

 0

 0

 0

 0

 23

 0

 0

 0

 0

 0

 0

 0

 0

 0

 0

 0

 0

 0

 0

 0

 0

 0

 0

 0

 0

 0

 0

 0

 0

 0

 0

 0

 0

 0

 0

 0

 0

 0

 0

 0

 0

 0

 0

 0

 0

 0

 0

 0

 0

 0

 0

 0

 0

 0

 0

 0

 0

 0

 0

 0

 0

 0

 0

 0

 0

 0

 0

 0

 0

 0

 0

 0

 0

 0

 0

 0

 0

 0

 0

 0

 0

 0

 0

 0

 0

 0

 0

 0

 0

 0

 0

 0

 0

 0

 0

 0

 0

 0

 0

 0

 0

 0

 0

 0

 0

 0

 0

 0

 0

 12

 12

 0

 0

 0

 0

 0

 0

 0

 0

 0

 0

 0

 0

 0

 0

 0

 0

 0

 0

 0

 0

 0

 0

 0

 0

 0

 0

 0

 0

 12

 3

 0

 0

 0

 0

 0

 0

 0

 0

 0

 0

 0

 0

 0

 0

 0

 0

 0

 0

 3

 0

 0

 0

 0

 0

 0

 0

 0

 0

 0

 0

 0

 0

 3

 0

 0

 0

 0

 0

 7

 0

 0

 0

 0

 0

 0

 0

 0

 0

 0

 0

 0

 0

 0

 0

 0

 0

 0

 0

 0

 0

 0

 0

 0

 0

 7

 7

 0

 0

 0

 0

 0

 0

 0

 0

 0

 0

 0

 0

 0

 1

 1

 1

 0

 0

 0

 0

 0

 0

 0

 0

 0

 0

 0

 0

 0

 0

 0

 0

 0

 0

 0

 0

 0

 0

 2

 0

 0

 0

 0

 0

 0

 0

 0

 0

 0

 0

 0

 0

 0

 0

 0

 0

 0

 0

 0

 0

 0

 0

 0

 0

 0

 0

 0

 0

 0

 0

 0

 0

 0

 2

 2

 0

 2

 0

 0

 0

 0

 0

 0

 0

 0

 0

 0

 0

 0

 0

 0

 0

 0

 0

 0

 0

 0

 0

 0

 0

 0

 0

 0

 5314

 70

 70

 70

 0

 0

 0

 17

 11

 6

 0

 0

 0

 1

 0

 4

 6

 2

 4

 1

 1

 1

 0

 0

 0

 131

 131

 0

 0

 131

 4950

 128

 128

 3312

 262

 0

 0

 0

 0

 0

 2

 189

 319

 0

 0

 0

 0

 0

 0

 0

 3

 0

 0

 2014

 63

 0

 460

 1505

 42

 436

 46

 0

 0

 0

 0

 0

 11

 929

 41

 0

 0

 5

 0

 0

 0

 5

 4

 0

 0

 0

 0

 0

 0

 0

 0

 0

 0

 4

 2

 2

 0

 0

 0

 0

 26

 2

 2

 0

 0

 0

 0

 0

 0

 13

 0

 0

 13

 0

 1

 1

 1

 0

 0

 1

 0

 0

 9

 9

 0

 0

 0

 0

 0

 0

 0

 3

 3

 3

 60

 10

 10

 0

 0

 0

 0

 2

 2

 0

 0

 0

 0

 0

 0

 0

 0

 0

 0

 0

 0

 0

 0

 0

 1

 1

 0

 0

 6

 6

 0

 0

 0

 0

 41

 41

 0

 0

 0

 24

 24

 24

 15

 15

 15

 13

 0

 0

 12

 10

 0

 0

 0

 0

 2

 1

 0

 0

 1

 0

 0

 0

 0

 103

 103

 0

 0

 2

 0

 2

 0

 0

 101

 1

 0

 49

 51

 0

 0

 0

 0

 0

 0

 0

 0

 807

 264

 264

 264

 0

 0

 0

 0

 0

 0

 0

 0

 0

 0

 0

 18

 0

 0

 18

 18

 0

 0

 0

 0

 0

 0

 0

 0

 0

 0

 0

 0

 0

 0

 0

 0

 0

 0

 0

 0

 0

 0

 0

 0

 0

 0

 0

 0

 0

 0

 0

 0

 0

 0

 0

 0

 0

 0

 0

 0

 3

 3

 3

 0

 0

 0

 0

 0

 0

 18

 18

 18

 42

 0

 0

 0

 0

 0

 0

 0

 40

 0

 40

 0

 0

 0

 0

 0

 0

 0

 2

 2

 3

 3

 3

 0

 0

 0

 0

 0

 0

 0

 0

 0

 0

 0

 0

 0

 0

 0

 0

 0

 79

 79

 2

 0

 42

 2

 33

 0

 0

 0

 0

 0

 0

 0

 0

 2

 2

 0

 2

 0

 0

 0

 0

 0

 0

 0

 0

 26

 11

 11

 0

 0

 0

 0

 9

 9

 6

 6

 0

 0

 0

 0

 0

 0

 0

 0

 0

 0

 0

 0

 0

 0

 0

 0

 0

 0

 0

 0

 0

 0

 0

 0

 0

 0

 0

 0

 0

 0

 0

 0

 0

 0

 0

 0

 0

 0

 0

 0

 0

 0

 0

 0

 0

 0

 0

 0

 0

 0

 0

 0

 0

 0

 0

 0

 0

 0

 0

 0

 0

 0

 0

 0

 0

 0

 0

 0

 0

 0

 0

 0

 0

 0

 0

 0

 0

 0

 0

 0

 0

 20

 20

 20

 0

 0

 0

 0

 0

 0

 0

 0

 0

 0

 0

 0

 0

 0

 0

 0

 0

 0

 0

 6

 6

 6

 0

 0

 0

 305

 0

 0

 0

 0

 0

 0

 288

 2

 0

 2

 0

 0

 0

 283

 1

 17

 3

 0

 0

 0

 0

 0

 0

 0

 0

 14

 0

 0

 0

 0

 0

 0

 0

 0

 0

 0

 0

 0

 0

 0

 21

 2

 2

 19

 19

 0

 0

 0

 0

 0

 0

 0

 0

 0

 0

 0

 0

 0

 0

 0

 0

 0

 0

 8

 8

 8

 8

 0

 0

 0

 0

 0

 0

 0

 0

 0

 0

 0

 0

 0

 0

 0

 0

 0

 0

 0

 0

 0

 0

 0

 0

 0

 0

 0

 0

 0

 0

 0

 0

 0

 0

 0

 0

 0

 7

 7

 7

 7

 7

 0

 0

 0

 0

 0

 391

 391

 57

 57

 57

 12

 12

 12

 0

 0

 0

 0

 0

 0

 16

 16

 16

 0

 0

 0

 306

 0

 0

 0

 0

 0

 0

 0

 0

 0

 3

 3

 303

 0

 296

 0

 0

 7

 0

 0

 90

 90

 90

 90

 90

 9

 9

 9

 9

 9

 4

 4

 0

 0

 0

 0

 0

 0

 0

 0

 0

 0

 0

 0

 4

 4

 4

 0

 0

 0

 0

 0

 0

 0

 0

 0

 0

 4

 0

 0

 0

 0

 0

 0

 0

 0

 0

 0

 0

 0

 0

 0

 0

 3

 0

 0

 0

 1

 1

 1

 0

 0

 2

 2

 0

 0

 0

 0

 2

 0

 0

 0

 0

 0

 0

 0

 0

 0

 0

 0

 0

 0

 0

 0

 0

 0

 0

 0

 0

 0

 0

 0

 0

 0

 0

 1

 1

 0

 0

 0

 0

 0

 0

 0

 0

 1

 0

 0

 1

 0

 0

 0

 0

 0

 0

 0

 0

 0

 0

 0

 0

 0

 0

 0

 0

 0

 0

 0

 0

 0

 0

 0

 0

 0

 0

 0

 0

 0

 0

 0

 0

 0

 0

 0

 0

 0

 0

 0

 0

 0

 0

 0

 0

 0

 0

 0

 0

 0

 0

 1877

 1877

 1877

 1877

 1877

 1877
