## Supplemental Figure 4 for "Experimentally-validated correlation analysis reveals new anaerobic methane oxidation partnerships with consortium-level heterogeneity in diazotrophy": Supplemental_Figure_4.html

Javascript must be enabled to view this page.

magnitude

 20238

 13049

 0

 0

 0

 0

 0

 0

 0

 0

 0

 0

 0

 0

 0

 0

 0

 0

 0

 0

 13034

 0

 0

 0

 0

 0

 0

 0

 0

 4

 4

 0

 0

 0

 0

 1

 1

 0

 0

 3

 3

 0

 0

 0

 0

 0

 0

 0

 0

 0

 0

 0

 0

 13028

 0

 0

 0

 0

 0

 0

 0

 0

 0

 0

 0

 0

 0

 0

 0

 0

 0

 0

 0

 0

 0

 0

 0

 0

 0

 0

 0

 0

 0

 0

 13028

 3

 3

 12994

 82

 12550

 362

 15

 15

 0

 0

 0

 0

 0

 0

 16

 16

 0

 0

 0

 0

 0

 0

 0

 0

 0

 0

 0

 2

 0

 0

 0

 0

 0

 0

 0

 0

 0

 1

 1

 1

 0

 0

 0

 0

 0

 0

 1

 0

 0

 0

 0

 0

 0

 0

 0

 0

 0

 1

 1

 0

 0

 0

 0

 0

 0

 0

 0

 0

 0

 0

 0

 0

 0

 0

 0

 0

 0

 0

 0

 0

 0

 0

 0

 0

 0

 0

 0

 0

 15

 0

 0

 0

 0

 0

 0

 0

 0

 0

 0

 0

 0

 0

 0

 0

 0

 4

 4

 4

 4

 0

 0

 0

 0

 10

 10

 10

 10

 1

 0

 0

 0

 1

 1

 0

 0

 1

 0

 0

 0

 0

 0

 0

 0

 0

 0

 0

 0

 0

 0

 0

 0

 0

 0

 0

 0

 0

 0

 0

 0

 0

 0

 0

 0

 0

 0

 0

 0

 0

 0

 0

 0

 0

 0

 6562

 19

 19

 19

 19

 19

 646

 1

 1

 1

 1

 7

 0

 0

 0

 0

 0

 0

 0

 0

 0

 0

 0

 0

 0

 0

 0

 0

 0

 0

 0

 0

 0

 1

 1

 1

 0

 0

 0

 0

 0

 0

 2

 2

 2

 0

 0

 0

 0

 0

 0

 0

 0

 0

 0

 0

 0

 0

 0

 0

 0

 0

 0

 0

 0

 0

 0

 0

 0

 0

 0

 0

 0

 0

 0

 2

 2

 2

 2

 2

 2

 634

 0

 0

 0

 0

 0

 0

 0

 0

 0

 0

 0

 0

 0

 0

 0

 0

 7

 0

 0

 0

 0

 0

 0

 0

 0

 7

 7

 0

 0

 627

 627

 627

 0

 0

 0

 0

 0

 0

 4

 4

 4

 4

 0

 0

 0

 0

 32

 0

 0

 0

 0

 22

 22

 0

 0

 0

 0

 0

 0

 0

 0

 22

 22

 0

 0

 0

 0

 0

 0

 6

 0

 0

 0

 0

 0

 0

 0

 0

 0

 0

 0

 0

 0

 0

 0

 0

 0

 0

 0

 0

 0

 0

 0

 0

 0

 0

 0

 0

 0

 0

 0

 2

 0

 0

 0

 0

 0

 0

 2

 2

 0

 0

 0

 0

 0

 0

 0

 0

 0

 0

 0

 0

 0

 0

 0

 0

 0

 0

 0

 4

 0

 0

 0

 0

 4

 0

 0

 1

 3

 0

 0

 0

 0

 0

 0

 0

 0

 0

 0

 0

 0

 0

 0

 0

 0

 0

 0

 0

 0

 3

 3

 3

 3

 0

 0

 0

 0

 0

 0

 0

 0

 0

 0

 0

 0

 0

 0

 0

 0

 1

 1

 1

 1

 0

 0

 0

 0

 0

 0

 0

 0

 0

 0

 0

 0

 0

 0

 0

 0

 0

 0

 0

 0

 0

 0

 0

 0

 0

 0

 0

 0

 0

 0

 0

 0

 0

 0

 0

 0

 0

 0

 0

 0

 0

 0

 0

 0

 0

 0

 0

 0

 0

 0

 0

 0

 0

 0

 0

 0

 0

 810

 102

 102

 102

 102

 127

 127

 127

 127

 0

 0

 0

 0

 0

 0

 0

 0

 13

 13

 0

 0

 0

 0

 0

 0

 13

 0

 0

 13

 0

 0

 0

 0

 0

 0

 0

 0

 0

 0

 0

 16

 16

 0

 0

 0

 0

 0

 0

 0

 0

 0

 0

 0

 0

 0

 16

 1

 0

 0

 0

 0

 0

 0

 0

 0

 6

 0

 9

 0

 0

 0

 0

 0

 0

 0

 233

 233

 0

 0

 0

 0

 17

 3

 0

 0

 0

 0

 0

 14

 216

 190

 3

 0

 0

 0

 0

 0

 0

 0

 0

 0

 5

 0

 0

 0

 0

 0

 0

 0

 0

 0

 0

 0

 0

 0

 3

 15

 0

 0

 0

 0

 0

 0

 0

 0

 0

 0

 7

 7

 7

 7

 76

 76

 76

 76

 50

 50

 1

 1

 0

 0

 0

 0

 0

 0

 0

 0

 0

 0

 0

 0

 0

 0

 0

 0

 0

 0

 0

 0

 0

 0

 1

 0

 0

 0

 1

 0

 0

 0

 0

 0

 0

 48

 48

 0

 0

 186

 186

 186

 186

 0

 0

 0

 0

 0

 0

 0

 0

 0

 0

 0

 0

 0

 0

 0

 0

 0

 0

 0

 0

 7

 7

 7

 7

 7

 13

 13

 13

 13

 13

 0

 0

 0

 0

 0

 0

 0

 0

 0

 0

 0

 0

 0

 0

 0

 0

 0

 0

 0

 0

 0

 0

 0

 0

 0

 0

 0

 0

 43

 43

 43

 43

 43

 0

 0

 0

 0

 0

 0

 0

 0

 0

 0

 310

 0

 0

 0

 0

 0

 0

 0

 0

 310

 310

 310

 310

 0

 0

 0

 0

 0

 0

 0

 0

 0

 0

 0

 0

 0

 0

 0

 0

 0

 0

 97

 0

 0

 0

 0

 0

 0

 97

 97

 11

 11

 1

 1

 0

 0

 0

 0

 82

 82

 3

 3

 269

 3

 3

 3

 3

 250

 250

 250

 1

 0

 0

 0

 0

 249

 6

 0

 0

 0

 6

 6

 6

 2

 2

 2

 0

 2

 0

 0

 0

 0

 0

 0

 0

 0

 0

 0

 0

 0

 0

 0

 0

 0

 0

 0

 0

 0

 0

 0

 0

 0

 0

 0

 0

 0

 0

 0

 0

 0

 0

 0

 0

 0

 0

 0

 0

 0

 0

 0

 0

 0

 0

 0

 0

 0

 0

 0

 0

 0

 0

 0

 0

 0

 0

 0

 0

 0

 0

 0

 0

 0

 0

 0

 0

 0

 0

 0

 0

 0

 0

 0

 0

 0

 0

 0

 0

 0

 0

 0

 0

 0

 0

 0

 0

 0

 0

 0

 0

 0

 0

 0

 0

 0

 0

 0

 0

 0

 0

 0

 0

 0

 0

 8

 8

 8

 8

 6

 0

 0

 0

 0

 0

 0

 0

 0

 6

 0

 0

 0

 6

 6

 0

 0

 0

 0

 6

 0

 0

 0

 0

 0

 0

 0

 0

 0

 0

 0

 0

 0

 0

 0

 0

 0

 0

 0

 0

 0

 0

 0

 0

 0

 0

 0

 15

 15

 15

 0

 0

 0

 0

 10

 10

 0

 0

 0

 0

 5

 5

 0

 0

 0

 0

 0

 0

 0

 0

 0

 0

 0

 0

 0

 0

 0

 0

 0

 0

 1

 1

 0

 0

 0

 0

 0

 0

 0

 0

 0

 0

 0

 0

 0

 0

 0

 1

 1

 1

 0

 0

 0

 0

 0

 0

 0

 0

 0

 0

 0

 0

 0

 0

 0

 2

 2

 0

 0

 0

 0

 0

 0

 0

 0

 0

 0

 0

 0

 0

 0

 0

 0

 0

 0

 0

 0

 0

 0

 0

 2

 2

 2

 99

 0

 0

 0

 0

 2

 0

 0

 0

 2

 0

 0

 0

 0

 0

 2

 0

 2

 0

 0

 0

 0

 0

 0

 0

 0

 0

 0

 0

 0

 0

 0

 0

 0

 0

 0

 0

 0

 0

 0

 0

 0

 0

 0

 0

 0

 0

 0

 0

 0

 0

 0

 0

 0

 0

 0

 0

 0

 0

 0

 0

 0

 0

 0

 0

 0

 97

 0

 0

 0

 97

 4

 4

 2

 2

 23

 0

 0

 23

 13

 9

 0

 0

 0

 4

 0

 0

 0

 0

 0

 0

 0

 0

 0

 0

 0

 0

 0

 0

 0

 0

 0

 0

 0

 0

 0

 0

 1

 0

 1

 0

 0

 0

 0

 0

 0

 0

 0

 0

 0

 0

 0

 0

 0

 0

 0

 2

 0

 0

 0

 0

 2

 0

 0

 4

 4

 2

 0

 0

 0

 0

 0

 0

 2

 28

 0

 15

 0

 13

 16

 1

 0

 0

 0

 15

 0

 0

 0

 0

 0

 0

 0

 0

 0

 0

 2

 2

 0

 0

 0

 0

 0

 0

 0

 0

 0

 0

 0

 0

 0

 0

 0

 0

 0

 0

 0

 0

 0

 0

 0

 0

 0

 0

 0

 0

 0

 0

 0

 0

 0

 0

 0

 0

 0

 0

 0

 0

 0

 0

 0

 0

 0

 0

 0

 0

 0

 0

 0

 0

 30

 30

 0

 0

 0

 4

 4

 4

 2

 2

 2

 0

 0

 0

 24

 24

 24

 0

 0

 0

 16

 16

 16

 16

 16

 0

 0

 0

 0

 0

 0

 0

 0

 0

 0

 51

 3

 3

 3

 3

 0

 0

 0

 0

 0

 0

 0

 0

 0

 0

 0

 0

 0

 0

 0

 0

 0

 0

 0

 0

 0

 0

 0

 0

 0

 0

 0

 0

 0

 0

 0

 0

 0

 0

 0

 1

 1

 1

 1

 3

 0

 0

 0

 0

 0

 0

 3

 3

 3

 0

 0

 0

 0

 38

 38

 38

 38

 0

 0

 0

 0

 0

 0

 0

 0

 4

 4

 4

 4

 0

 0

 0

 0

 2

 2

 2

 2

 1

 1

 1

 1

 1

 1

 1

 1

 0

 0

 0

 0

 1

 1

 0

 0

 0

 0

 0

 0

 0

 0

 0

 0

 0

 0

 0

 0

 0

 0

 0

 0

 0

 86

 0

 0

 0

 0

 0

 0

 0

 0

 0

 0

 0

 0

 0

 0

 0

 0

 0

 0

 0

 0

 6

 6

 6

 6

 24

 1

 1

 1

 0

 0

 0

 0

 0

 0

 0

 0

 0

 0

 0

 0

 0

 0

 0

 0

 0

 0

 0

 0

 0

 3

 3

 3

 14

 14

 14

 0

 0

 0

 6

 3

 3

 0

 0

 0

 0

 1

 1

 2

 0

 0

 2

 0

 0

 0

 0

 0

 0

 0

 0

 0

 0

 0

 0

 0

 0

 0

 0

 0

 0

 0

 0

 0

 0

 0

 0

 0

 0

 0

 0

 0

 0

 0

 0

 2

 2

 2

 2

 54

 0

 0

 0

 0

 0

 54

 54

 3

 3

 0

 0

 0

 0

 21

 0

 2

 2

 0

 0

 23

 0

 0

 0

 0

 0

 0

 0

 0

 0

 0

 0

 0

 3943

 12

 12

 12

 12

 0

 0

 0

 0

 0

 0

 0

 0

 0

 0

 0

 0

 25

 0

 0

 0

 0

 0

 0

 0

 0

 0

 0

 0

 0

 0

 0

 0

 0

 0

 0

 0

 0

 0

 0

 0

 0

 0

 0

 0

 0

 0

 0

 0

 0

 0

 0

 0

 6

 0

 0

 0

 0

 0

 0

 0

 0

 0

 0

 0

 0

 0

 0

 0

 0

 0

 0

 3

 2

 0

 0

 0

 0

 0

 0

 1

 0

 0

 0

 0

 0

 0

 0

 0

 0

 0

 2

 0

 0

 0

 0

 0

 0

 2

 0

 0

 0

 0

 0

 0

 0

 0

 0

 1

 0

 0

 0

 0

 1

 0

 0

 0

 0

 0

 0

 0

 0

 16

 16

 1

 0

 0

 0

 0

 2

 0

 0

 0

 0

 0

 0

 0

 0

 0

 0

 0

 0

 0

 0

 0

 0

 0

 0

 0

 1

 0

 0

 12

 1

 0

 0

 0

 0

 0

 0

 0

 0

 0

 0

 0

 0

 0

 0

 0

 0

 0

 0

 1

 0

 0

 0

 0

 0

 0

 0

 0

 0

 0

 0

 0

 0

 1

 0

 0

 0

 0

 0

 2

 0

 0

 0

 0

 0

 0

 0

 0

 0

 0

 2

 2

 0

 0

 0

 0

 0

 0

 0

 0

 0

 0

 0

 0

 0

 0

 0

 0

 0

 0

 0

 0

 0

 0

 0

 0

 0

 0

 0

 0

 0

 0

 0

 0

 0

 0

 0

 0

 0

 0

 0

 0

 0

 0

 0

 0

 0

 0

 0

 0

 0

 0

 0

 0

 0

 0

 0

 0

 0

 0

 0

 0

 0

 0

 0

 0

 0

 0

 0

 0

 0

 0

 0

 0

 0

 0

 0

 0

 0

 0

 0

 0

 0

 0

 0

 0

 0

 0

 0

 0

 0

 0

 0

 0

 0

 0

 0

 0

 0

 0

 0

 0

 0

 0

 0

 0

 0

 0

 0

 0

 0

 0

 0

 0

 0

 0

 0

 0

 0

 0

 2418

 20

 20

 20

 0

 0

 0

 0

 0

 0

 0

 0

 0

 0

 0

 0

 0

 0

 0

 0

 0

 0

 0

 0

 0

 95

 95

 0

 0

 95

 2256

 16

 16

 1533

 52

 0

 0

 0

 0

 0

 0

 15

 1397

 0

 0

 0

 0

 0

 0

 0

 0

 0

 0

 16

 13

 0

 40

 707

 123

 87

 271

 2

 0

 0

 0

 0

 2

 0

 220

 0

 2

 0

 0

 0

 0

 0

 0

 0

 0

 0

 0

 0

 0

 0

 0

 0

 0

 0

 0

 0

 0

 0

 0

 0

 16

 3

 3

 0

 0

 0

 0

 0

 0

 4

 0

 0

 4

 0

 1

 1

 0

 0

 0

 0

 0

 0

 8

 8

 0

 0

 0

 0

 0

 0

 0

 0

 0

 0

 15

 0

 0

 0

 0

 0

 0

 0

 0

 0

 0

 0

 0

 0

 0

 0

 0

 0

 0

 0

 0

 0

 0

 0

 5

 1

 0

 4

 1

 1

 0

 0

 0

 0

 9

 9

 2

 2

 2

 3

 3

 3

 9

 9

 9

 2

 0

 0

 1

 0

 0

 0

 0

 0

 1

 1

 0

 0

 1

 0

 0

 0

 0

 789

 789

 0

 0

 0

 0

 0

 0

 0

 789

 2

 0

 146

 641

 0

 0

 0

 0

 0

 0

 0

 0

 693

 65

 65

 65

 0

 0

 0

 0

 0

 0

 0

 0

 0

 0

 0

 3

 0

 0

 2

 2

 0

 0

 0

 0

 0

 0

 0

 0

 0

 0

 0

 0

 0

 0

 0

 0

 0

 0

 0

 0

 0

 0

 0

 0

 0

 0

 0

 0

 0

 0

 0

 0

 0

 1

 1

 0

 0

 0

 0

 0

 0

 0

 0

 0

 0

 0

 0

 0

 0

 19

 19

 19

 96

 0

 0

 1

 0

 1

 0

 0

 95

 0

 92

 0

 0

 3

 0

 0

 0

 0

 0

 0

 6

 6

 6

 0

 0

 0

 1

 1

 1

 0

 0

 0

 0

 0

 0

 0

 0

 0

 0

 0

 1

 1

 0

 0

 0

 0

 1

 0

 0

 0

 0

 0

 0

 0

 0

 0

 0

 0

 0

 0

 0

 0

 0

 0

 0

 0

 0

 10

 7

 7

 0

 0

 0

 0

 3

 3

 0

 0

 0

 0

 0

 0

 0

 0

 0

 0

 0

 0

 0

 0

 0

 0

 0

 0

 0

 0

 0

 0

 0

 0

 0

 0

 0

 0

 0

 0

 0

 0

 0

 0

 0

 0

 0

 0

 0

 0

 0

 0

 0

 0

 0

 0

 0

 0

 0

 0

 0

 0

 0

 0

 0

 0

 0

 0

 0

 0

 0

 0

 0

 0

 0

 0

 0

 0

 0

 0

 0

 0

 0

 0

 0

 0

 0

 0

 0

 0

 0

 0

 0

 389

 389

 389

 0

 0

 0

 0

 0

 0

 0

 0

 0

 0

 0

 0

 0

 0

 0

 0

 0

 0

 0

 9

 9

 9

 0

 0

 0

 70

 0

 0

 0

 0

 0

 0

 56

 9

 0

 2

 0

 0

 0

 43

 2

 14

 1

 0

 0

 0

 0

 2

 0

 0

 0

 11

 0

 0

 0

 0

 0

 0

 0

 0

 0

 0

 0

 0

 0

 0

 24

 0

 0

 22

 22

 0

 0

 0

 0

 0

 2

 2

 0

 0

 0

 0

 0

 0

 0

 0

 0

 0

 0

 4

 4

 4

 4

 0

 0

 0

 0

 0

 0

 0

 0

 0

 0

 0

 0

 0

 0

 0

 0

 0

 0

 0

 0

 2

 2

 2

 2

 0

 0

 0

 0

 0

 0

 0

 0

 0

 0

 0

 0

 0

 1

 1

 1

 1

 1

 0

 0

 0

 0

 0

 25

 25

 0

 0

 0

 3

 3

 3

 0

 0

 0

 0

 0

 0

 1

 1

 1

 0

 0

 0

 21

 0

 0

 0

 0

 0

 0

 0

 0

 0

 0

 0

 21

 0

 19

 0

 0

 2

 0

 0

 5

 5

 5

 5

 5

 8

 8

 8

 8

 8

 3

 3

 0

 0

 0

 2

 2

 2

 0

 0

 0

 0

 0

 0

 1

 1

 1

 0

 0

 0

 0

 0

 0

 0

 0

 0

 0

 23

 0

 0

 0

 0

 0

 0

 0

 0

 0

 0

 0

 0

 0

 0

 0

 2

 0

 0

 0

 2

 2

 2

 0

 0

 0

 0

 0

 0

 0

 0

 0

 0

 0

 0

 0

 0

 0

 0

 0

 0

 0

 0

 0

 0

 0

 0

 0

 0

 0

 0

 0

 0

 0

 0

 0

 0

 0

 21

 21

 4

 4

 0

 0

 1

 1

 0

 0

 16

 0

 0

 15

 1

 0

 0

 0

 0

 0

 0

 0

 0

 0

 0

 0

 0

 0

 0

 0

 0

 0

 0

 0

 0

 0

 0

 0

 0

 0

 0

 0

 0

 0

 0

 0

 0

 0

 0

 0

 0

 0

 0

 0

 0

 0

 0

 0

 0

 0

 0

 0

 0

 0

 627

 627

 627

 627

 627

 627
